# The Translation Initiation Factor Homolog, *eif4e1c*, Regulates Cardiomyocyte Metabolism and Proliferation During Heart Regeneration

**DOI:** 10.1101/2022.08.15.502524

**Authors:** Anupama Rao, Baken Lyu, Ishrat Jahan, Anna Lubertozzi, Gao Zhou, Frank Tedeschi, Eckhard Jankowsky, Junsu Kang, Bryan Carstens, Ken Poss, Kedryn Baskin, Joseph Aaron Goldman

## Abstract

The eIF4E family of translation initiation factors bind 5’ methylated caps and act as the limiting-step for mRNA translation. The canonical eIF4E1A is required for cell viability, yet other related eIF4E families exist and are utilized in specific contexts or tissues. Here, we describe a family called Eif4e1c for which we find roles during heart development and regeneration in zebrafish. The Eif4e1c family is present in all aquatic vertebrates but is lost in all terrestrial species. A core group of amino acids shared over 500 million years of evolution forms an interface along the protein surface, suggesting Eif4e1c functions in a novel pathway. Deletion of *eif4e1c* in zebrafish caused growth deficits and impaired survival in juveniles. Mutants surviving to adulthood had fewer cardiomyocytes and reduced proliferative responses to cardiac injury. Ribosome profiling of mutant hearts demonstrated changes in translation efficiency of mRNA for genes known to regulate cardiomyocyte proliferation. Although *eif4e1c* is broadly expressed, its disruption had most notable impact on the heart and at juvenile stages. Our findings reveal context-dependent requirements for translation initiation regulators during heart regeneration.

## Introduction

Zebrafish regenerate heart muscle after catastrophic injury. Fervent research is underway to discover molecules that regulate fundamental events in regeneration and have potential relevance to regenerative medicine. Changes in the abundance of dozens to thousands of mRNAs have been reported using both targeted approaches like in-situ hybridization and unbiased approaches based on RNAseq (Goldman et al., 2017; Honkoop et al., 2018; Wu et al., 2016). However, reliance on mRNA as the sole measure of gene expression is an imperfect assumption at best, with only ∼40% of mRNA levels correlating with the amount of protein (Buccitelli and Selbach, 2020). Similarly, the abundance of mRNA and proteins are poorly interconnected during heart regeneration, suggesting an important role for post-transcriptional regulation (Ma et al., 2018). An alternative snapshot of expression levels was revealed by profiling of mRNA bound to a transgenic ribosomal subunit (Fang et al., 2013), which uncovered a role for JAK/STAT signaling in heart regeneration. However, few total transcripts were identified, likely due to ribosome heterogeneities (Emmott et al., 2018; Genuth and Barna, 2018), the dependence on transgenesis, and/or limited starting material. Regulation of translation is an important mechanism that impacts gene expression (Kong and Lasko, 2012; Wang et al., 2020a) and is a central feature of physiologic heart growth (Chorghade et al., 2017). How such regulation might impact heart regeneration, however, remains largely unexplored until now.

Pathways that initiate translation are diverse, yet the classical means is through loading of the ribosome onto start codons via recognition of 5’ methylated caps (Borden and Volpon, 2020). The canonical eIF4E1A major cap binding protein is the rate-limiting factor to initiate loading, and its expression levels help determine the translation of specific mRNA cohorts (Davis et al., 2019; Truitt et al., 2015). In mouse, eIF4E1A, the canonical *Eif4e1a* gene, is essential for development and viability (Sénéchal et al., 2021) limiting most mechanistic studies of its function to cell culture. However, studies of partial knockdowns and heterozygotes have shown *Eif4e1a* is generally found in excess and at reduced levels improves resiliency to cancer in heterozygous mice (Graff et al., 2007; Truitt et al., 2015) and indicated improved aging in adult worms (Syntichaki et al., 2007). Actively growing heart muscle in neonatal mice has increased eIF4E1A levels (Chen et al., 2022), but how eIF4E1A is regulated in a context of innate heart regeneration is unknown. While the canonical eIF4E1A promotes translation, a second family of eIF4E1 homologs called eIF4E1B has been reported to have repressive functions (Minshall et al., 2007; Robalino et al., 2004). In fish, a third close relative of canonical eIF4E1A, called *eif4e1c*, has been reported (Joshi et al., 2005; Zhang et al., 2018), but its function and properties have not yet been described.

In a previous profiling study, we found that expression of the *eif4e1c* gene increases in zebrafish hearts during regeneration (Goldman et al., 2017). We show that *eif4e1c* represents a new and highly conserved family of eIF4E1 that is retained in aquatic vertebrate and lost in terrestrial animals. The evolutionary changes that maintain heart regeneration in some organisms and not others are an area of intense study but with questions that still remain (Hirose et al., 2019; Lai et al., 2017; Stockdale et al., 2018; Wang et al., 2020b). Here, using CRISPR generated mutants, we demonstrate that fish-specific *eif4e1c* is required for normal production of cardiomyocytes in zebrafish hearts during animal growth and injury-induced regeneration. Furthermore, our findings indicate that regulation of gene expression through translational mechanisms is a participating component of cardiogenesis.

## Results and Discussion

### Eif4e1c is a family of translation initiation factors conserved in all aquatic vertebrates

Zebrafish have two paralogs of the canonical eIF4E1A, *eif4ea* and *eif4eb*, that are likely a result of a whole genome duplication in teleost fish ∼380 million years ago (Taylor et al., 2001). Unlike the canonical eIF4E1A, zebrafish have only single copies of *eif4e1b* and *eif4e1c* (Joshi et al., 2005) raising the possibility that they are duplicates of one another. Since the eIF4E1A and eIF4E1B families have distinct roles in translation, the evolutionary history of *eif4e1c* may help inform its function. To understand the relationship of *eif4e1c* to the two other eIF4E1 families, we undertook a phylogenetic analysis of eukaryotic eIF4E1 orthologs. Using the IQ-TREE web server (Trifinopoulos et al., 2016) we conducted a maximum likelihood analysis to estimate the phylogeny of 142 homologs from 53 species ranging from yeast to humans (Fig. S1A). Interestingly, Eif4E1c forms its own clade that is ancestral to the split of the Eif4e1a and Eif4e1b families (Fig. 1). A homolog of *eif4e1c* was found in all 37 aquatic vertebrates from this sampling including non-teleosts, suggesting that the Eif4e1c family predates the teleost specific duplication event. Whereas the Eif4e1a and Eif4e1b families are shared by aquatic and terrestrial vertebrate species, the Eif4e1c clade is absent from terrestrial animals. We conclude that the Eif4e1c family must have been lost in an early ancestor of terrestrial vertebrates when coelacanth split from amphibians and is retained in all aquatic vertebrate fishes.

**Fig. 1.**
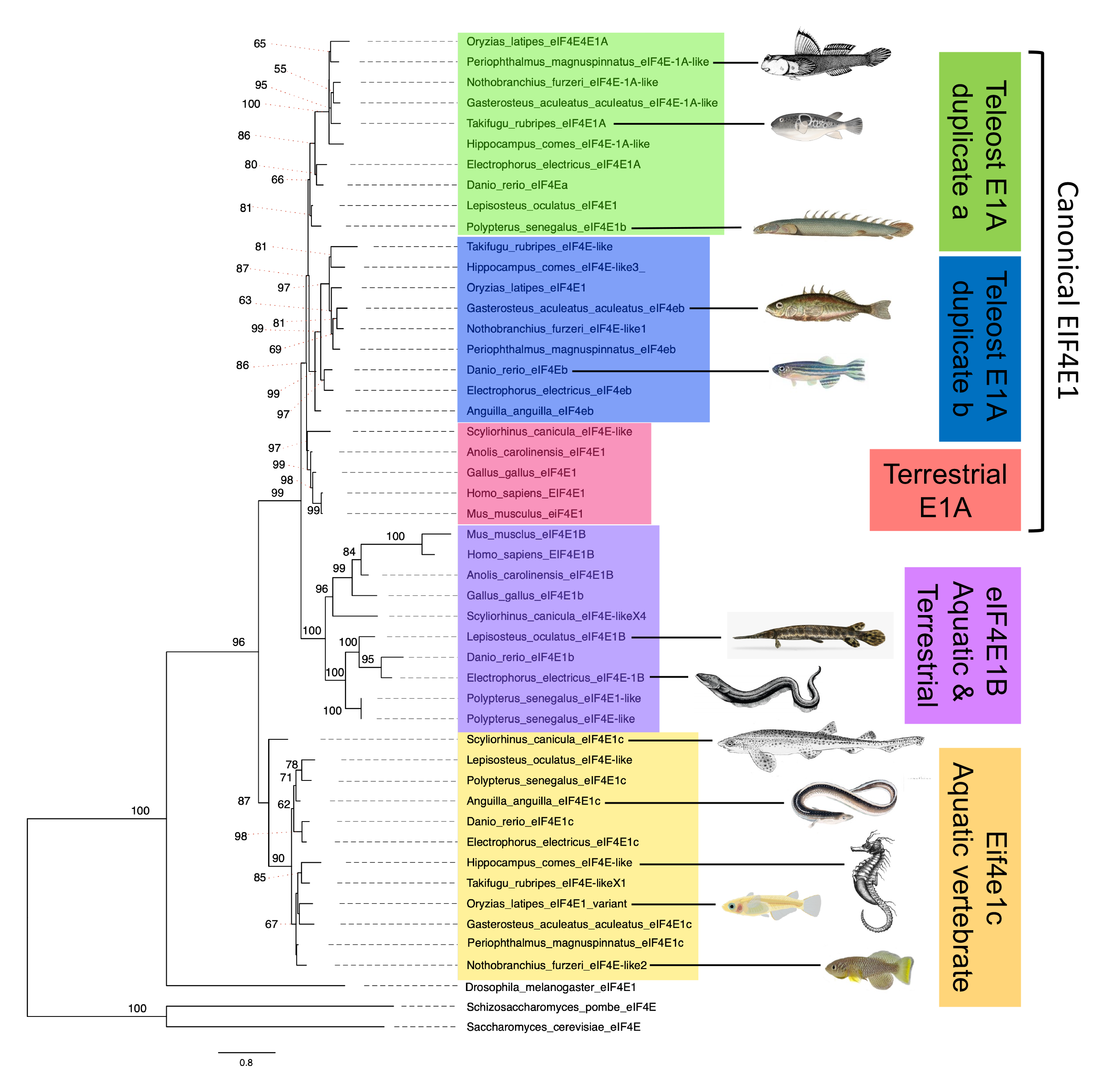
Eif4e1c is both unique to and shared by all aquatic vertebrates. Shown is a phylogeny of eIF4E1 orthologues from a sampling of the species considered (see Fig. S1 for full analysis). Eif4e1c is ancestral to the canonical eIF4E1A split from its variant, eIF4E1B. Terrestrial species have a canonical and eIF4E1B ortholog (pink and purple). All aquatic species have a canonical variant: 8 species with a duplication (blue and green), 2 species retain only one variant of the duplication, and the *Scyliorhinus canicular* (Small-spotted catshark) canonical clusters with terrestrial eIF4E1A. All 12 aquatic vertebrate retain an Eif4e1c family member. Shown are images of each of the aquatic vertebrate species to highlight the diversity considered. Only 5 of 12 aquatic vertebrate shown here retain an eIF4E1B variant. Estimates are made using maximum likelihood and the IQ-TREE. Nodes with bootstrap support < 0.85 are marked with their respective values, all other nodes had support values of 0.85 or higher.

To identify the sequence features that distinguish the Eif4e1c family we performed ClustalW alignment between the four zebrafish homologs and their human orthologs (Fig. 2A). All six homologs share the essential amino acids for binding 5’ methylated caps (green) and assembly of the preinitiation complex with eIF4G (pink). In the 12 amino acids reported to distinguish eIF4E1A and eIF4E1B families (Kubacka et al., 2015), *eif4e1c* has substantially more similarity to eIF4E1A. For example, there are 7 residues identical or similar to canonical eIF4E1A (gray boxes) and 4 residues identical to the eIF4E1B variant (black boxes); however, of the 6 residues that confer optimal binding affinity for caps (red carrots), all 6 in *eif4e1c* share similarity with the canonical eIF4E1A. ClustalW alignment within the Eif4e1c family showed that the evolutionary conservation is striking (Fig. S1B). Shark and zebrafish Eif4e1c are 86% identical and 95% similar with only 8/186 substantial amino acid differences in the protein core over ∼500 million years of evolution. There are 23 amino acids that are invariant in almost all Eif4e1c family members (Fig. 2A, yellow boxes). Most of the 23 Eif4e1c residues are at locations in the protein where changes have been found and tolerated in other homologs from isolated species. Interestingly, 7 of the 23 amino acids remain identical in *all* eIF4E1A and eIF4E1B orthologs throughout evolution but are only different in the Eif4e1c family (red boxes). Thus, from amino acid sequence comparisons, we conclude that while Eif4e1c family shares features with the canonical eIF4E1A in critical cap binding residues, Eif4e1c is likely a unique homolog with both highly conserved and distinctive properties.

**Fig. 2.**
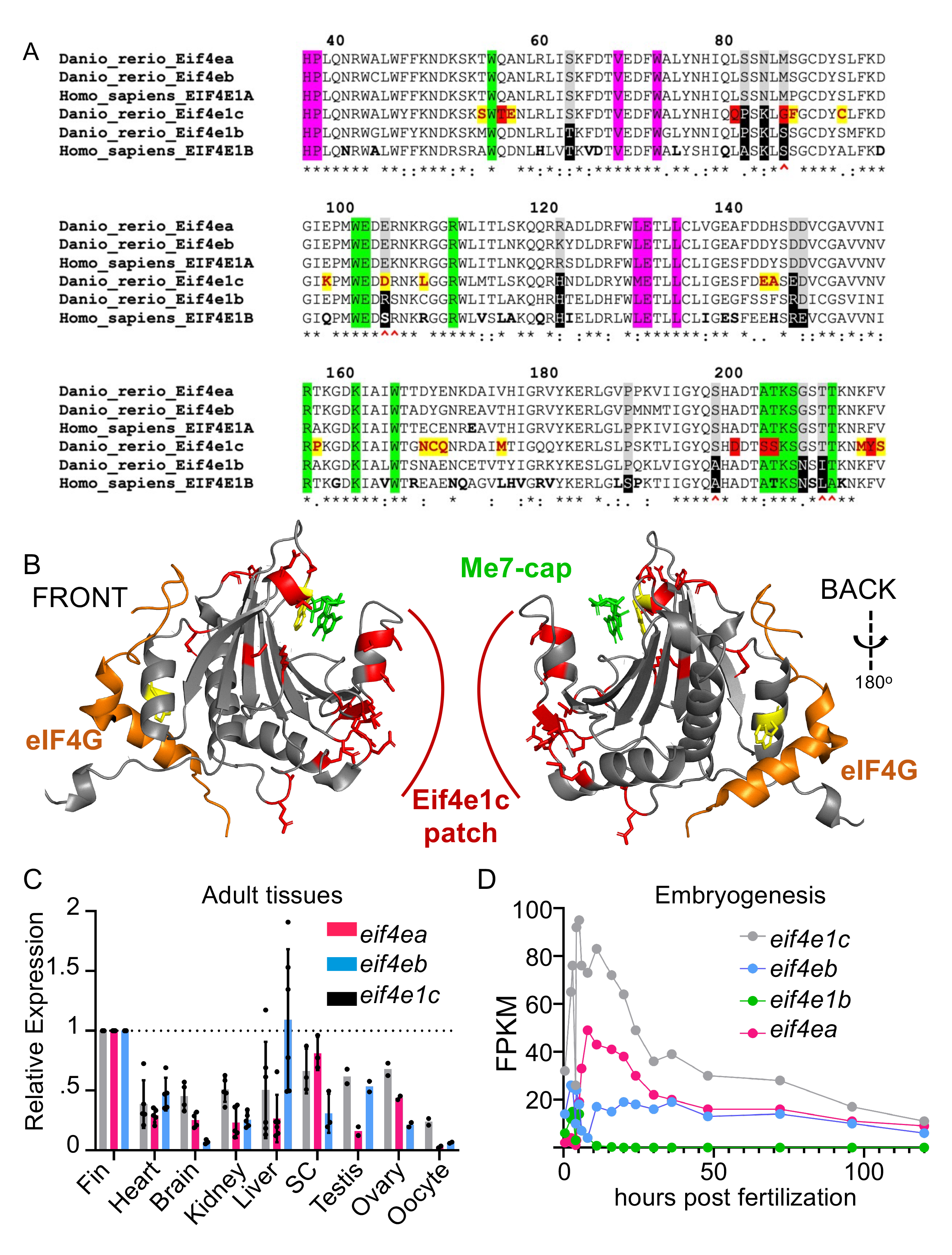
Highly conserved Eif4e1c specific amino acids form a novel patch along the surface of the protein. (A) ClustalW sequence alignment of human vs fish eif4E orthologs. Highlighted are universally conserved amino acids required for cap binding (green) and eIF4G recruitment (pink). Residues that distinguish EIF4E1A family members (gray) from EIF4E1B family members (black) are also highlighted. The six amino acids most responsible for strong cap binding are marked by red carrots (**^**). The Eif4e1c family conserved residues are highlighted yellow with red letters. The seven residues that are identical in all EIF4E1A and EIF4E1B family orthologs throughout evolution but are uniquely changed in Eif4e1c family members are highlighted in red. Amino acids that diverge between mouse and human are bolded in the human sequence. (B) Shown is a PyMol rendition of zebrafish Eif4e1c mapped along the human structure of EIF4E1A bound to a methylated cap (green) and a EIF4G peptide (orange). The tryptophans required for both associations are conserved and highlighted in yellow. All Eif4e1c specific residues are highlighted in red. (C) The abundance of each of the transcripts shown in the legend were derived from RNA sequencing data sets from different adult tissues. Expression is determined relative to *mob4.* and. The fpkm for the three orthologs were similar in the fin graphed values were then normalized to expression levels in the fin Where error bars are presented, at least three biological replicates were used to determine the average and standard deviation. Only the testis, ovaries, and oocyte had two replicates. Dots show individual data points. (D) Shown is a time course of RNA sequencing of different developmental stages in embryos. Normalized sequencing reads for the four different EIF4E1 orthologs are shown.

Crystal structures of eIF4E1A have been solved for several species showing all homologs retain conserved features. To understand how the Eif4e1c distinctive residues may impact properties of the protein, we modeled zebrafish Eif4e1c over a crystal structure of human EIF4E1A bound to a capped GTP residue and a peptide from EIF4G (Grüner et al., 2016) using RoseTTA fold (Fig. 2B) (Baek et al., 2021). The canonical folds of the modeled zebrafish Eif4e1c protein are identical to what is found in the core of all eIF4E1 family members, with similar positioning of the two highly conserved tryptophans required for cap binding and preinitiation complex assembly (yellow). This mirrors what is seen in the Alphafold prediction of the two molecules (Jumper et al., 2021). The 23 amino acids characteristic of the Eif4e1c family are positioned mainly along the protein surface in solvent exposed regions (Fig. 2B, red). A distinctive Eif4e1c specific patch is found opposite the EIF4G binding site and adjacent to the cap-binding region. Therefore, the highly conserved amino acids shared by the Eif4e1c family create a novel binding interface on the protein, suggesting that Eif4e1c interacts with unique partners.

### The *eif4e1c* gene is widely expressed in tissues and throughout development

Canonical eIF4E1A and eIF4E1B are known to have distinct expression patterns. For example, *eif4e1b* is only expressed in the gonads and skeletal muscle in adults and during the first 12 hours-post-fertilization (hpf) (Robalino et al., 2004). To determine if *eif4e1c* is expressed in all tissues like eIF4E1A or is restricted to discrete organs or developmental stages like *eif4e1b*, we surveyed expression of *eif4e1c* in published RNAseq data sets (see Methods). Each of *eif4ea, eif4eb,* and *eif4e1c* genes were expressed in every organ examined (Fig. 2C). Expression of *eif4e1c* was highest in the fin and in other tissues ranged from 38% to 67% of that total. These differences are likely within the typical RNAseq variability due to batch effects, so we conclude that *eif4e1c* is widely expressed at similar levels throughout zebrafish organ systems. While *eif4e1c* expression is widespread, whether it is confined to a particular cell-types within organs is unclear. We analyzed a published scRNAseq data set produced from adult zebrafish hearts that uncovered 15 different identifiable cell-types (Hu et al., 2022). Transcripts for *eif4e1c,* and the canonical *eif4ea* and *eif4eb*, were detected within each of the 15 clusters suggesting that all three transcripts are expressed in all cardiac cell-types including in CMs (Fig. S2). Thus, like its canonical orthologs, *eif4e1c* is broadly expressed in all organ systems and cell-types examined.

The canonical homologs *eif4ea* and *eif4eb* are broadly expressed at similar levels as *eif4e1c* except for low expression of *eif4eb* in the brain (6.5-fold decrease) and *eif4ea* in the testis (3.8- fold decreased). Interestingly, eif4e1c was found to be the predominant homolog present in oocytes, 8.5-fold higher than *eif4ea* and 3.7-fold higher than *eif4eb* (Fig. 2C). High levels of maternal deposition suggest that *eif4e1c* may play a critical role during embryogenesis. Using the EMBL expression atlas, we assessed expression levels of all eIF4E1 orthologs during a time course across early zebrafish development (White et al., 2017). Zebrafish *eif4e1c* is the most highly expressed eIF4E1 at every time point during the first five days of development, during which most organ systems are formed (Fig. 2D). Expression levels of *eif4e1c* are roughly equal to the sum of both *eif4ea* and *eif4eb* except during the first 12hpf where *eif4e1c* is 3-4X more abundant. During rapid growth stages such as during embryogenesis, *eif4e1c* is the dominantly expressed cap-binding homolog but is expressed mostly at similar levels as the canonical factors in the adult.

### Deletion of the *eif4e1c* locus causes poor growth and survival to adulthood

*Eif4e1a* in mice is required for viability with no detectable homozygous mutant mice from heterozygous crosses (Sénéchal et al., 2021). To examine whether *eif4e1c* is essential in the zebrafish system, we generated a stable CRISPR mutant fish line with an *eif4e1c* deletion, hereafter referred to as Δ*eif4e1c*. Using guide RNA targeting flanking loci, we created a 12,131bp deletion in the *eif4e1c* gene extending from intron 1 to exon 7 and deleting the entire coding region (Fig. 3A). Crosses between Δ*eif4e1c* heterozygotes yielded larvae at 3 days-post-fertilization (dpf) in normal Mendelian ratios (Fig. S3A). Based on the gene expression analysis above (Fig. 2D) it is possible that embryonic Δ*eif4e1c* mutant fish develop through early stages due to maternal deposition of wildtype *eif4e1c* mRNA or protein (Harvey et al., 2013). To address maternal deposition effects, we crossed homozygous Δ*eif4e1c* mutant females with heterozygous Δ*eif4e1c* males and still found Mendelian ratios in the progeny, eliminating maternal deposition as a requirement for Δ*eif4e1c* mutant fish development (58+/- : 49-/-). Mutants were found at normal Mendelian ratios through early juvenile stages up to 4 weeks-post-fertilization (wpf) (chi^2^ = 0.629, p value = 0.73). Beginning in late juvenile stages, 8wpf, heterozygotes and homozygous mutants were underrepresented among the progeny (599+/+ : 951-/+ : 318-/-, chi^2^ = 29.3, p value = 4.31 x 10^-7^). Although homozygous Δ*eif4e1c* fish were recovered in adults at 3 months-post-fertilization (mpf), survivorship was diminished in mutants and heterozygotes (Fig. S3A, chi^2^ = 84.9, p value = 3.22 x 10^-19^). Therefore, deletion of *eif4e1c* is not embryonically lethal but begins to influence viability between 4 and 8wpf with only ∼80% of heterozygotes and ∼50% of homozygote mutants surviving past 8wpf.

**Figure 3.**
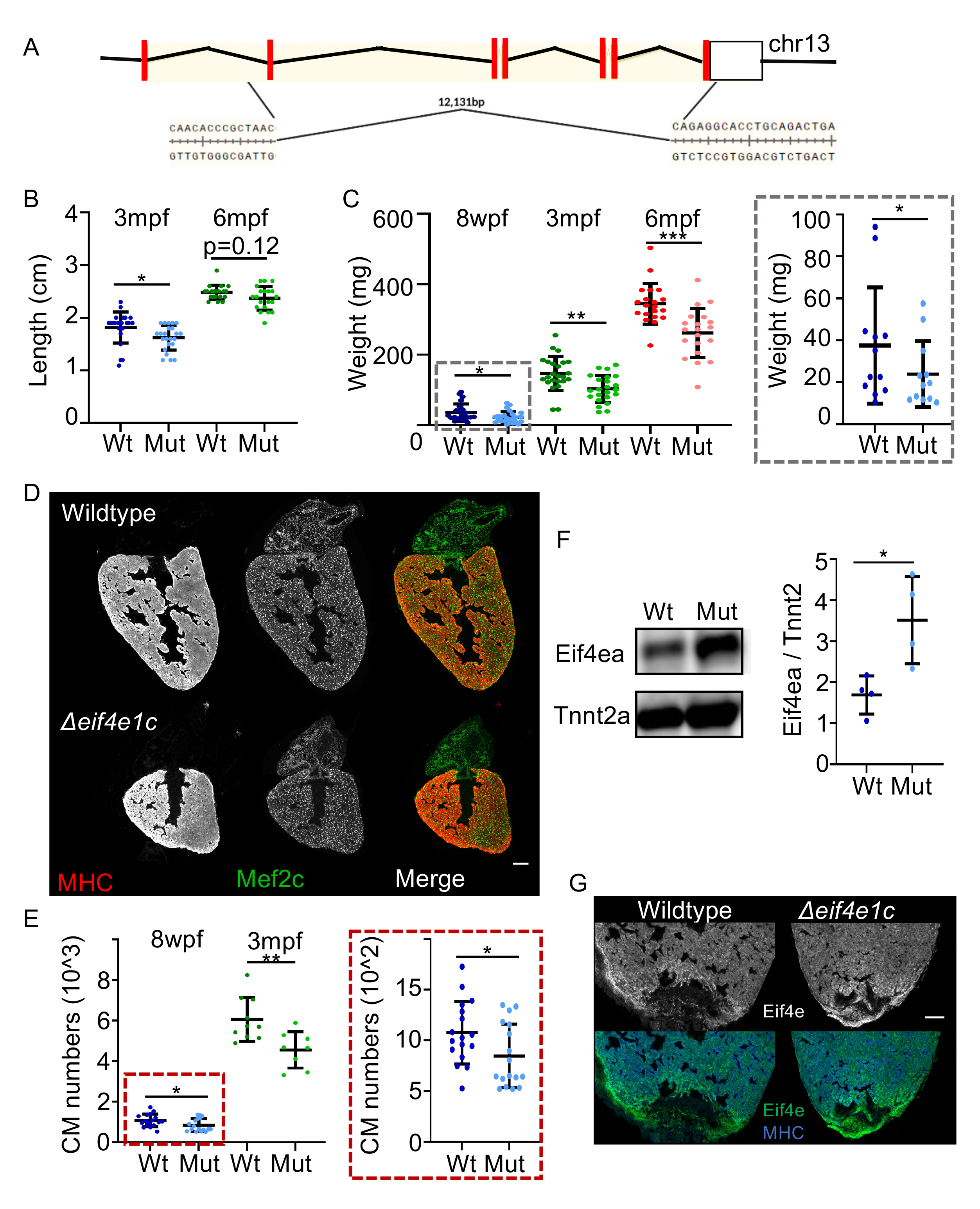
Δ*eif4e1c* mutants have impaired growth and poor survival. (A) Schematic of the CRISPR mediated deletion of *eif4e1c*. Red rectangles represent coding exons, and the white box represents exons with untranslated regions. (B) Fish were grown together, genotyped at the listed stage (top label) and immediately measured from jaw to the caudal fin bifurcation. (3- month average wildtype = 1.82cm and mutant = 1.62cm; Welch’s t-test, p-value = 0.0156, N = 24 vs 23; 6-month average wildtype = 2.48cm and mutant = 2.37cm; Mann-Whitney, p-value = 0.121, N = 20 vs 20). (C) After measuring length, fish were dried off as much as possible and weighed. Gray box is a blow-up of the 8wpf data with a different scale for the y-axis (8-week (blue) average wildtype = 35.75mg and mutant = 22.82mg; Welch’s t-test, p-value = 0.0392, N = 24 vs 24; 3-month (green) average wildtype = 146.5mg and mutant = 103.1mg; Welch’s t-test, p-value = 0.0014, N = 24 vs 23; 6-month average (red) wildtype = 344.6mg and mutant = 261.7mg; Welch’s t-test, p-value = 0.0002, N = 20 vs 20). (D) Uninjured heart ventricles from wildtype and mutant fish were sectioned and stained for muscle using antibodies toward the Myosin heavy chain (MHC) and the cardiac transcription factor Mef2c to identify CM nuclei (Scale bar = 100μm). (E)The numbers of Mef2c positive cells were counted with MIPAR. Gray box is a blow-up of the 8wpf data with a different scale for the y-axis. (8-week (blue) average wildtype= 1078 and mutant = 848, Welch’s t-test p-value = 0.0383, N = 17 vs 17; 3-month (green) average wildtype= 6061 and mutant = 4557; Welch’s t-test, p-value = 0.0052, N = 10 vs 8). (F) left - Western blot of whole cell extracts from zebrafish hearts using an antibody directed towards canonical Eif4ea (top) and Tnnt2a (bottom). Right – Quantification of western blots with total Eif4ea levels normalized to sarcomeric protein Tnnt2a as a measure of total cardiac mass (average increase = 1.82; Welch’s t-test, p-value = 0.0334, N = 4 vs 4). (G) Immunofluorescence of canonical Eif4ea/b (top – grayscale, bottom – green) and MHC (blue)(Scale bar = 100μm). For each panel, the lighter shade color is used for mutants, individual data points are represented by dots, and horizontal black bars display the mean (middle) or standard error (top and bottom).

Throughout all stages of development Δ*eif4e1c* mutants appear visibly normal morphologically but Δ*eif4e1c* mutant adults have notable size differences suggesting impaired growth. At 3mpf, Δ*eif4e1c* mutant fish were 12% shorter (Fig. 3B; mean length:wildtype= 1.816cm, mutant = 1.622cm, p-value= 0.0156, n=24 vs 23) with 30% lower mass (Fig. 3C; mean weight: wildtype = 146.5mg, mutant = 103.1mg, p-value= 0.0014, n=24, 23). Similar results were found when males and females were considered separately and heterozygous Δ*eif4e1c* carriers did not demonstrate significant length or weight differences (Fig. S3B). At 8wpf, Δ*eif4e1c* mutant fish had 36% less mass (Fig. 3C; mean weight: wildtype = 35.75mg, mutant = 22.82mg, p-value=0.0392, n=24, 24) and at 6mpf Δ*eif4e1c* mutant fish had 24% less mass (Fig. 3C; mean weight: wildtype = 344.6mg, mutant = 261.7mg, p-value= 0.0002, n= 20, 20). Therefore, growth differences between Δ*eif4e1c* mutant fish and wildtype fish peak at juvenile stages and decrease into adulthood (Fig. S3C). Similarly, length differences found at 3mpf are not detectable by 6mpf (Fig. 3B; mean length wildtype = 2.480cm, mutant = 2.370cm, p-value = 0.1212, Mann-Whitney, n= 20, 20). We conclude that Δ*eif4e1c* homozygous fish have reduced growth that persists but attenuates into adulthood.

To determine if apparent growth defects reflect fewer overall cell numbers, we quantified CM numbers using an antibody for a nuclear marker of cardiac muscle (Mef2c). CM content was compared between adult hearts (3mpf) of Δ*eif4e1c* mutant fish and their wildtype siblings. Mutant heart ventricles had 25% fewer CMs compared to their WT siblings at 3mpf (Fig. 3E; mean: wildtype = 6061, mutant = 4557, p-value = 0.0052, n=10, 8). Fewer CMs were also measured at 8wpf just after mutant death occurred (Fig. 3E, mean: wildtype = 1078, mutant = 848, p-value = 0.0388, n=17, 17). At 6mpf, mutant heart ventricles had fewer CMs demonstrating that fewer CMs during development are not fully rescued by regeneration mechanisms in the adult (Fig. S3D). Reduced CM numbers may result from less CM proliferation, reduced CM survival, or increased CM death. To see if hearts from Δ*eif4e1c* mutant fish were undergoing increased apoptosis, we performed TUNEL on heart sections from mutant and wildtype fish and observed no significant change (Fig. S3E-F). We conclude that deficits in overall growth of adult *eif4e1c* mutants likely reflect defects in either cell proliferation or cell survival during development. Phalloidin staining of sarcomeres revealed no gross differences between CM sarcomere structure in wildtype and mutant hearts from (Fig. S3G). Likely, proliferation or survival deficits are unrelated to structural differences in Δ*eif4e1c* mutant CMs.

In contrast to reported *Eif4e1a* mutants in mice, zebrafish *eif4e1c* deletion knockouts can survive (Altmann et al., 1989; Sénéchal et al., 2021). Similarities in sequence and expression patterns between *eif4e1c* and its canonical homologs *eif4ea* and *eif4eb* raises the possibility that Δ*eif4e1c* mutants survive because the canonical homologs can functionally substitute. To look at canonical protein levels, we used an antibody raised against the human canonical EIF4E1 that also recognizes the zebrafish orthologs (see Methods for details). Western blot of whole cell extracts from wildtype and Δ*eif4e1c* mutant hearts showed that canonical Eif4ea/b protein levels are increased in Δ*eif4e1c* mutants (Fig. 3F, fold change avg. = 1.82, N = 4,4). Interestingly, we see Eif4ea/b protein levels increase at the site of injury during wildtype heart regeneration (Fig. 3G). In both wildtype and Δ*eif4e1c* mutant hearts, after amputation of the apex of the ventricle, Eif4ea/b protein levels increase to a similar extent at the site of injury (Fig. S3H). Taken together, canonical eIF4E1 protein levels increase in Δ*eif4e1c* mutant hearts, as they do during wildtype heart regeneration. We conclude that canonical Eif4ea/b likely partially compensates for cardiac growth deficits in surviving Δ*eif4e1c* mutant hearts.

Studies in other organisms have shown a wide array of lower expression levels of canonical eIF4E1A (50-70%) without obvious phenotypes in growth (Graff et al., 2007; Truitt et al., 2015). Therefore, we do not believe that reduced total levels of *eif4e1a* paralogs underlie growth deficits in Δ*eif4e1c* mutant adults. These observations lead us to hypothesize that the *eif4e1c* ortholog is specialized and functions to foster growth in juveniles and adults, a phenomenon commonly observed in aquatic species. From the reduced body size of Δ*eif4e1c* mutants we speculate that loss of the Eif4e1c family in early terrestrial vertebrates may have limited growth in adults as an adaptation to survival on land.

### Ribosome profiling of Δ*eif4e1c* mutant hearts uncovers translational changes

The eIF4e1 family core amino acids responsible for initiating loading of ribosomes for translation are conserved in zebrafish Eif4e1c. Therefore, we hypothesized that the survival and growth deficits that occur upon deleting *eif4e1c* result from alterations in translation. To measure global translation in *eif4e1c* mutants we injected fish with O-propargyl-puromycin (OPP), which terminates peptide chain elongation. Total translation (incorporation of OPP) can be measured by fluorescence levels by conjugating fluorophores using CLIC chemistry (Fig. 4A). Injection of OPP into Δ*eif4e1c* mutants and wildtype siblings demonstrated that global protein synthesis is unperturbed in Δ*eif4e1c* mutants (Fig. 4B; mean fluorescence: wildtype = 140279 adu/sq.µm, mutant = 140596 adu/sq.µm; p-value = 0.806, Mann-Whitney, N = 15,15). We conclude that growth deficits in Δ*eif4e1c* mutants are not a result of impaired general translation.

**Figure 4.**
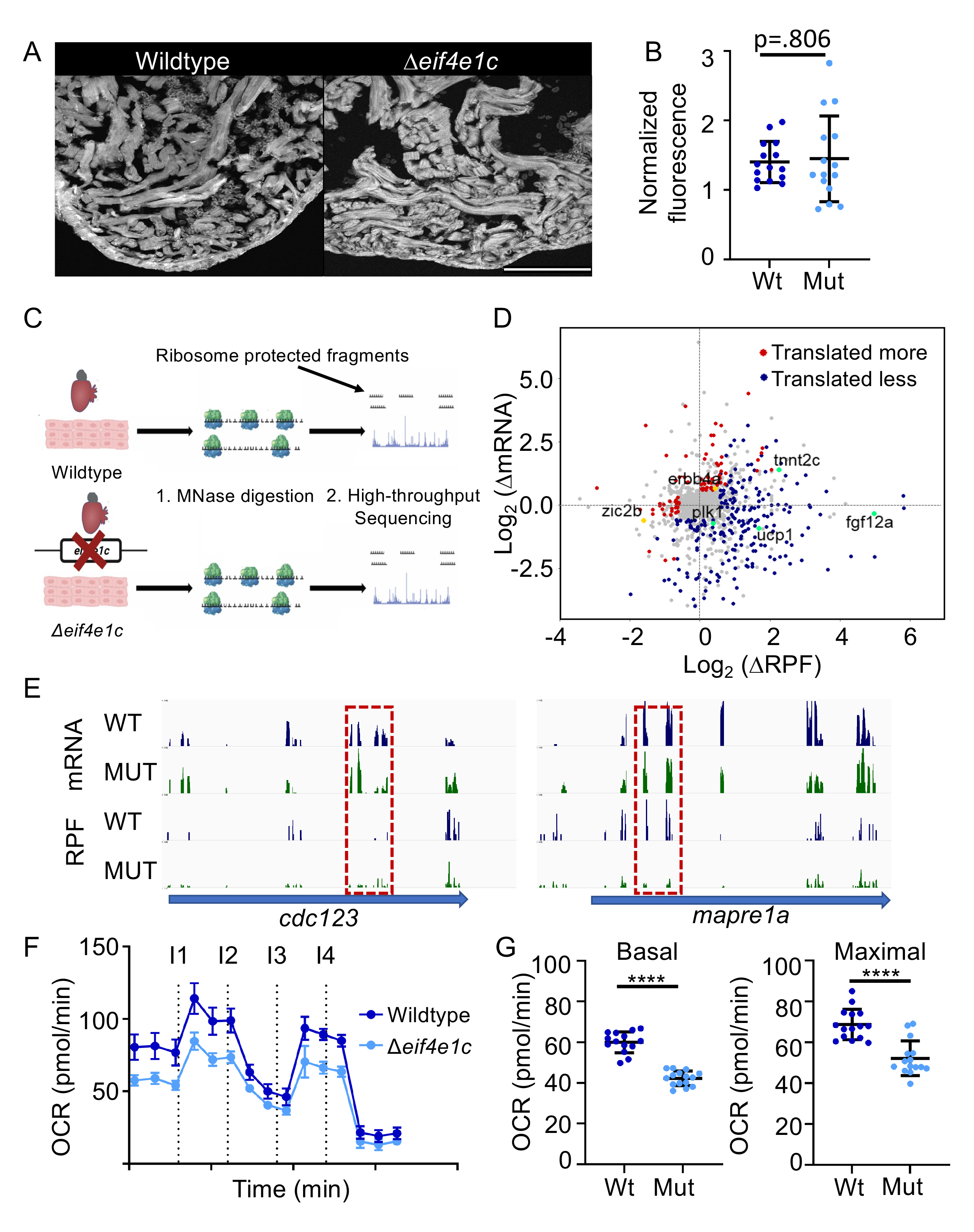
Ribosome profiling indicates translational changes in *eif4e1c* mutant hearts. (A) Shown is an example of fluorescence from hearts of OPP injected fish (Scale bar = 100μm). (B) Quantification of OPP fluorescence shows no significant difference between mutant and wildtype fish (normalized fluorescence avg. wildtype= 140279 adu/sq.µm and mutant = 145096 adu/sq.µm, Mann Whitney p-value = 0.806, N = 15 vs 15). Wildtype – Blue, Mutant – Light blue. Horizontal black bars display the mean (middle) or standard error (top and bottom). (C) Shown is a cartoon of ribosome profiling. Ribosome bound mRNA is purified from wildtype and mutant hearts and then subjected to micrococcal nuclease (MNase) digestion. Fragments of mRNA protected by being bound by ribosomes are then purified and subjected to high-throughput sequencing. (D) Differences (mutan/wildtype) in mRNA abundance (log_2_) are plotted versus changes in the abundance of ribosome protected fragments (RPF). Genes whose translation does not change significantly are plotted in gray, genes where translation is decreasing in the mutant are plotted in red and those genes where translation is increasing in the mutant are plotted in blue. Decreasing genes mentioned in the text are colored gold and increasing genes mentioned in the text are colored green. (E) Shown are high-throughput sequencing browser tracks for two genes (blue arrows) that are involved in cell-cycle progression (*cdc123*, and *mapre1a*). Wildtype (WT) data is shown in blue, and mutant (MUT) data is shown in green. The top two tracks (mRNA) show the abundance of mRNA (as measured from RNAseq) and the bottom two tracks show the abundance of ribosome protected fragments (RPF). Dashed red boxes highlight decreasing levels of RPF where mRNA levels are comparable. (F) Shown is a time-course of oxygen consumption rates (OCR) from mitochondria measured by the Seahorse analyzer. Hashed lines indicate time points of drug injection. Readings were normalized to the number of mitochondria using qPCR. The first injection (I1) is of ADP to stimulate respiration, the second injection (I2) is of oligomycin to inhibit ATP synthase (complex V) decreasing electron flow through transport chain, the third injection (I3) is of FCCP to uncouple the proton gradient, and the fourth injection (I4) is of antimycin A to inhibit complex III to shut down mitochondrial respiration. Wildtype – Blue, Mutant – Light blue. Horizontal black bars display the mean (middle) or standard error (top and bottom). (G) Basal respiration is calculated as the OCR average after addition of ADP (I1 to I2) subtracting the non-mitochondrial respiration after injection of antimycin A (after I4) (mean wildtype = 60.01 and mutant = 42.25; Welch’s t-test, p value < 0.0001; N = 15 vs 15). Maximal respiration is calculated as the OCR average after FCCP addition (I3 to I4) subtracting the non-mitochondrial respiration after injection of antimycin A (after I4) (mean wildtype = 68.73 and mutant = 52.19; Welch’s t-test, p value < 0.0001; N = 15 vs 15). Wildtype – Blue, Mutant – Light blue. Horizontal black bars display the mean (middle) or standard error (top and bottom).

To determine if particular transcripts are translated differently in *eif4e1c* mutants we performed ribosome profiling which involves sequencing mRNA fragments protected by virtue of being bound to the ribosome (Fig. 4C). Therefore, the level of ribosome protected fragments (RPF) can illuminate the translational status of a transcript. Normalization of RPFs to the total amount of transcript (RNAseq) provides a quantitative measure of translational efficiency that can be compared between conditions. Since, Δ*eif4e1c* mutants have small hearts as adults, we performed ribosome profiling to compare the translatome of adult 3mpf Δ*eif4e1c* mutant hearts to that of their wildtype siblings. Overall, there were 88 genes with transcripts that are less efficiently translated and 224 genes with transcripts that were more efficiently translated in the Δ*eif4e1c* mutant (Fig. 4D). Thus, deletion of the *eif4e1c* gene resulted in translational changes in about one fifth (22.6%) of the total expressed genes detected in ribosome profiling of adult hearts.

Several of the translationally misregulated transcripts in Δ*eif4e1c* mutant hearts could explain some of the observed phenotypes. For example, RPF for *erbb4,* a receptor for the potent CM mitogen Nrg1, are 44% lower in Δ*eif4e1c* mutant hearts (Bersell et al., 2009; D’Uva et al., 2015; Gemberling et al., 2015). Retinoic acid signaling is also required for CM proliferation during development (Keegan et al., 2005) and regeneration (Kikuchi et al., 2011) The translational efficiency for a transcription factor downstream of retinoic acid signaling called *zic2b* is 1.6-fold lower in mutant hearts (Fig. 4D). In addition, several cell cycle genes had transcripts not translated as well in Δ*eif4e1c* mutant hearts (Fig. 4E). For example, translated less are *mapre1a* that associates with the AuroraB kinase to assemble microtubules during cytokinesis (Sun et al., 2008) and *cdc123* encodes a factor needed for protein synthesis initiation and cell cycle progression (Okuda and Kimura, 1996; Perzlmaier et al., 2013). We cannot formally determine from which cell-types these translational changes are occurring. Yet, we predict at least some changes are occurring in CMs since CMs compromise most cells in uninjured zebrafish hearts and decreased translation of cell proliferation transcripts like *mapre1a* and *cdc123* are likely occurring there. In sum, deletion of *eif4e1c* led to decreased translation of mRNA encoding signaling pathways that promote CM proliferation and cell division in uninjured hearts.

Other *eif4e1c*-dependent changes in translation may occur at time points closer to 4wpf when mutants first begin to die. Ribosome profiling at these stages is not possible since zebrafish hearts are too small to yield the amount of starting material required. By 12wpf translational buffering (Kusnadi et al., 2021; Lorent et al., 2019) or compensation from canonical cap-binding proteins (Fig. 3F) may mask some of the *eif4e1c* affected transcripts to support the 50% mutant survival beyond 4wpf. Evidence for compensation is present in the ribosome profiling data set. For example, we detected *increased* translational efficiency of transcripts encoding pro-growth factors in Δ*eif4e1c* mutant hearts such as Fgf family members (*fgf8b* - 2.96-fold, *fgf12a* - 31.59-fold, *fgf19* - 2.38-fold) that have been reported to be pro-survival and pro-proliferative to CMs (Lepilina et al., 2006; Sakurai et al., 2013; Tahara et al., 2021). In addition, reduced mitochondrial activity has been reported to be pro-proliferative in the heart (Cardoso et al., 2020; Fukuda et al., 2020; Honkoop et al., 2018; Miklas et al., 2021). Translated more efficiently are transcripts for *ucp1* (3.23-fold), a mitochondrial uncoupling factor that reduces mitochondrial activity to mitigate production of reactive oxygen species (Echtay et al., 2002). In conclusion, translation efficiency of different pro-growth pathways and molecules is both enhanced and diminished in Δ*eif4e1c* mutant hearts. Since mutant hearts are still deficient in growth (Fig. 3D-E), we postulate that *eif4e1c* is critical for heart growth but that other less efficient pro-growth pathways can partially compensate for *eif4e1c* loss through the canonical *eif4e1* orthologs.

To confirm the biological significance of increased *ucp1* translation we measured oxygen consumption rates (OCR) of mitochondria isolated from hearts using the Agilent Seahorse bioanalyzer Agilent (Fig. S4B). Mitochondria from Δ*eif4e1c* mutant hearts had lower OCR compared to the mitochondria from wildtype sibling hearts (Fig. 4F). The basal and maximal respiration rates of mutant hearts were significantly lower (25.59% and 24.06%, respectively), which suggests impaired mitochondrial output (Fig. 4G). To examine whether reduced mitochondrial activity is also observed by the Seahorse during wildtype heart growth, we compared respiration of mitochondria from uninjured hearts to mitochondria from regenerating hearts using the Seahorse analyzer. In agreement with a previous report (Honkoop et al., 2018), mitochondrial activity plummets in zebrafish hearts undergoing regeneration with basal respiration decreasing 75.5% and maximal respiration decreasing 62.0% (Fig. S4C-E). Since reduced mitochondrial activity correlates with growing heart muscle, a similar but less profound reduction in mitochondrial activity in Δ*eif4e1c* mutants may function as a mechanism to foster compensatory growth.

Ribosome profiling demonstrated that translational changes occur in Δ*eif4e1c* mutant hearts but it is unlikely every transcript is a direct Eif4e1c target. Second order effects are possible especially if compensation drives survival in Δ*eif4e1c* mutants. Therefore, it is likely some Eif4e1c-bound transcripts are not affected in our ribosome profiling experiment. Nevertheless, many translational changes *are* detected in hearts of Δ*eif4e1c* mutant hearts, including some that may explain their smaller heart size.

### Deletion of *eif4e1c* impairs metabolic remodeling and regeneration in the heart

We hypothesized that increased expression of *eif4e1c* mRNA in regenerating hearts may stimulate heart growth during regeneration (Goldman et al., 2017). After surgical removal of the apex of the heart, Δ*eif4e1c* mutants had diminished CM proliferation during the peak of heart regeneration compared to wildtype siblings (Fig. 5A). The fraction of Mef2c positive CMs that were also positive with the proliferation marker EdU was reduced by 35% seven days post amputation (dpa) of the ventricular apex (Fig. 5B, mean: wildtype = 10.68% and mutant = 6.021%, p-value = 0.0054, N = 11 vs 13). We cannot determine if *eif4e1c* proliferation phenotypes are CM specific or a result of secondary effects from other cell-types. Homozygous Δ*eif4e1c* mutants did complete regeneration despite early defects in CM proliferation and look the same as their wildtype siblings by 28dpa (Fig. S5A). Thus, we conclude that Δ*eif4e1c* mutants have impaired CM proliferation that may be resolved by compensatory mechanisms at later stages of regeneration.

**Figure 5.**
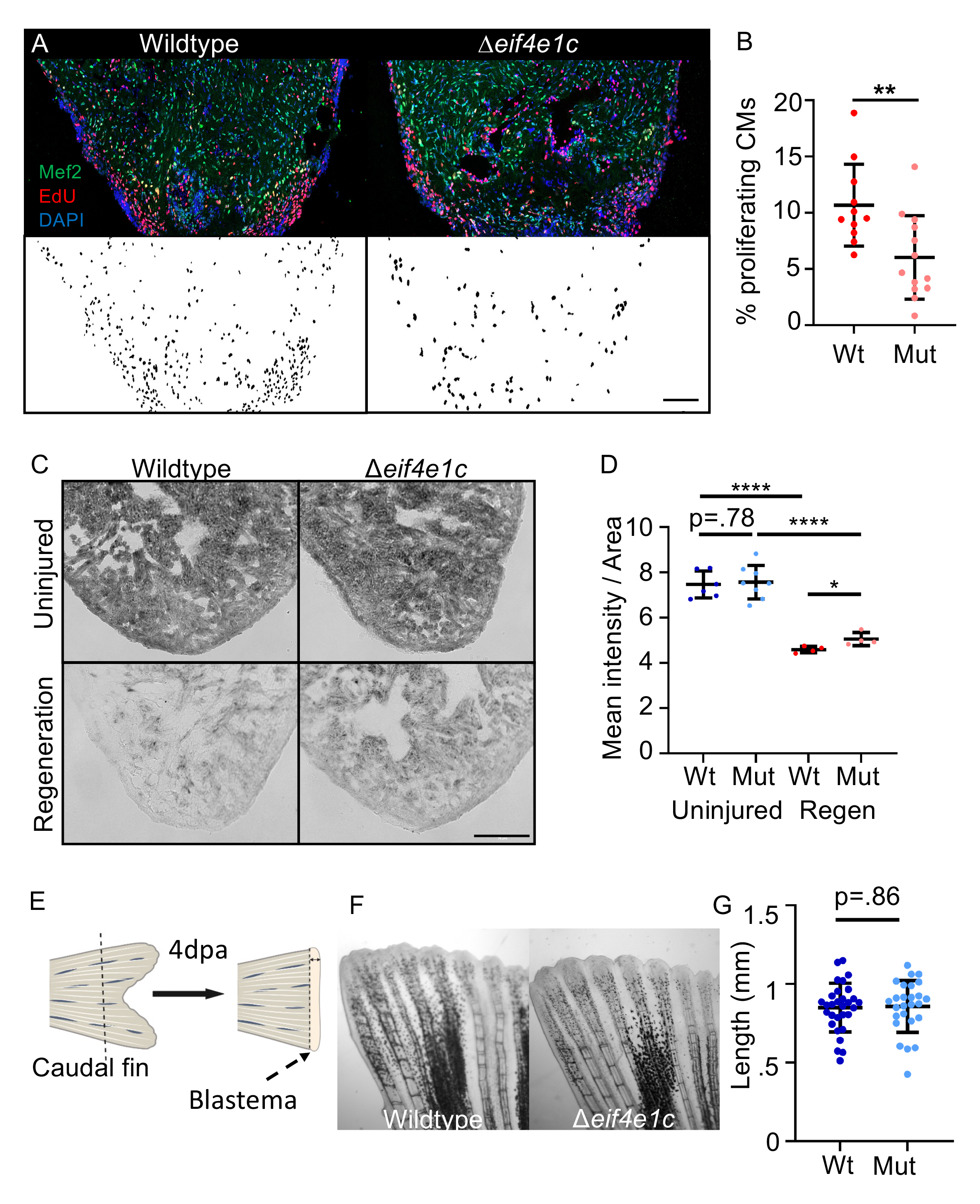
Deletion of *eif4e1c* impairs CM proliferation in regenerating hearts. (A) Images of sectioned amputated ventricles (7dpa) from wildtype and Δ*eif4e1c* mutant fish. Sections are stained for Mef2c (green) and EdU (red). Below – double positive cells are highlighted in black using a MIPAR software rendition (Scale bar = 100μm). (B) Quantification of CM proliferation indices (Mef2/EdU double positive over total Mef2 positive) in 7dpa ventricles (average = 10.68% and 6.02%; Welch’s t-test, **p-value = 0.0054, N = 11 vs 13). Wildtype – red, mutant – pink. Horizontal black bars display the mean (middle) or standard error (top and bottom). (C) Cryosections of fresh mutant and wildtype hearts were stained for Succinate dehydrogenase activity. Shown are uninjured hearts (top row) and hearts that were ablated genetically using ZCAT (7 days-post-incubation) (Scale bar = 20μm). (D) The mean signal intensity was calculated per given area for Sdh stained hearts (Uninjured average wildtype = 7.47 and mutant = 7.57 with N = 6 vs 8; Regeneration average wildtype = 4.59 and mutant = 5.06 with N = 4 vs 4). The Welch’s t-test was used to calculate significance (wildtype vs mutant uninjured = 0.781, wildtype uninjured vs wildtype regeneration **** < 0.0001, mutant uninjured vs mutant regeneration **** < 0.0001). For every replicate (N = 4 vs 4) the mutant Sdh activity during regeneration (pink) was higher than the wildtype activity during regeneration (red). Wildtype uninjured – blue, Mutant uninjured – light blue. Horizontal black bars display the mean (middle) or standard error (top and bottom). (E) Cartoon of caudal fin regeneration experiments. The blastema is the highly proliferative zone of growth that forms the new fin. (F) Caudal fins were amputated ∼50% in Δ*eif4e1c* mutants and their wildtype siblings and shown are images of fin regrowth 4 days later. (G) There was no significant difference between fin growth in wildtype (blue) and mutant (light blue) fish (average = 0.849mm and 0.857mm; Welch’s t-test, p-value = 0.861, N = 30 vs 26). Horizontal black bars display the mean (middle) or standard error (top and bottom).

As previously discussed, decreased mitochondrial respiration is a feature of heart regeneration that promotes growth. One of the key enzymes central to connecting the TCA cycle to the electron transport chain is succinate dehydrogenase (Sdh); Sdh activity is reported to be sharply downregulated during heart regeneration (Bezawork-Geleta et al., 2017; Honkoop et al., 2018). Using the <u>z</u>ebrafish <u>c</u>ardiomyocyte <u>a</u>blation transgenes (ZCAT), we injured CMs throughout the heart and confirmed that Sdh activity is sharply reduced during regeneration (Fig. 5C). While Δ*eif4e1c* mutant hearts also have lower Sdh activity during regeneration, Sdh activity was higher in Δ*eif4e1c* mutant hearts compared to their wildtype siblings (Fig. 5D). For uninjured hearts, Sdh activity was unchanged between mutant and wildtype. This suggests that an inability to fully reduce Sdh activity may in part underlie the impairment of heart regeneration in Δ*eif4e1c* mutants. It is unclear if the *sdh* transcript is a direct target of *eif4e1c* or if changes in its activity are regulated indirectly.

After development, Δ*eif4e1c* mutants had deficiencies in overall size (Fig. 3B-C). To determine whether impaired growth is a general feature of regeneration, we analyzed the ability of Δ*eif4e1c* mutants to regenerate their fins. Fin regeneration was determined by measuring growth of caudal fins in Δ*eif4e1c* mutants and their wildtype siblings 4dpa (Fig. 5E-F). There is no significant difference in fin regrowth indicating that Δ*eif4e1c* mutants can normally regenerate their fins (Fig. 5G; mean wildtype = 0.849 mm, mutant = 0.857 mm, p-value = 0.861, N = 30 vs 26). Earlier time points of fin regeneration at 2dpa similarly showed no difference between wildtype and Δ*eif4e1c* mutant siblings (Fig. S5B; mean: wildtype = 0.689 mm, mutant = 0.617 mm, p-value = 0.366, N = 18 vs 22). We performed a survey of published RNAseq data sets from zebrafish fins and found that while overall *eif4e1c* expression is highest in the uninjured fin (Fig. 2C), transcripts for *eif4e1c* are not increased during fin regeneration like they are in the heart (Fig. S5C-D, mean uninjured wildtype fin = 0.795, mean regenerating wildtype fin = 0.935, mean uninjured wildtype heart = 0.308, mean regenerating WT heart = 0.865). We conclude that while Δ*eif4e1c* mutants may have general growth deficits during development (Fig. 3BC), during regeneration *eif4e1c* function is more critical to growth in the heart.

In uninjured hearts and fins, transcript levels for *eif4e1c* are reduced 96% and 97% from wildtype levels in the mutants (Fig. S5E). As expected, the mutant transcript retains the 3’UTR but is missing most of the coding sequence and is therefore likely degraded. Interestingly, *eif4e1c* transcript levels during heart regeneration increase 10.7-fold in mutants which is nearly 5-fold higher than *eif4e1c* increases in the wildtype hearts during regeneration (Fig. S5F, mean: wildtype = 2.30, mutant = 10.70). This suggests a mechanism of compensation where a feedback loop stimulates *eif4e1c* transcription when Eif4e1c activity is absent. Such a feedback loop does not exist in regenerating fins since there was no difference in the change of *eif4e1c* transcript abundance between wildtype and mutants (Fig. S5F, mean: wildtype = 0.667, mutant = 0.787). The existence of a feedback loop in the heart and not in the fin further argues that Eif4e1c activity is more critical to the heart during regeneration. We cannot exclude that other compensation mechanisms may mask the presence of this feedback loop in fins.

This study establishes a new pathway for translational regulation mediated by a homolog of an mRNA cap-binding protein called *eif4e1c*. This pathway is highly conserved in aquatic species but lost in terrestrial vertebrates. Importantly, we show that in zebrafish, *eif4e1c* is required for normal cardiogenesis during development and regeneration. While *eif4e1c* is broadly expressed both during development and in adulthood, overt mutant phenotypes are restricted to particular tissues and first observed in juveniles. For example, heart regeneration, but not fin regeneration, is impaired. Likely, specific features of the Eif4e1c protein allow for targeting distinct cohorts of transcripts that underlie the different tissue sensitivities found during regeneration. The data presented here suggests that regulation of metabolism may be an aspect of *eif4e1c* function that predisposes heart muscle to a reliance of *eif4e1c*. Alternative pro-CM growth pathways not dependent on *eif4e1c* exist, as Δ*eif4e1c* mutants do complete regeneration albeit with a reduced index of CM proliferation. In the future, determining a contribution to regeneration for each of the canonical mRNA cap-binding proteins will help construct a translation regulatory network for heart regeneration with possible therapeutic implications.

## Materials and Methods

### Phylogeny of the Eif4e1c family

Expanding on previous studies, we analyzed 20 species of teleosts covering most if not all subclades (Lin et al., 2017) including the most ancestral species Asian arowana (*S. formosus*) (Bian et al., 2016) and the spotted gar (*L. oculatus*) (Braasch et al., 2016). Beyond teleost we also included multiple species of sharks, rays, eels, and the fish that are the closest relatives to terrestrial tetrapods like mudskippers (*P. magnuspinnatus*) (You et al., 2014) and lungfish (*P. senagalus*, *P. annectens*) (Austin et al., 2015). Homologs for *eif4e1c* were not found in more primitive deuterostomes, for example the sea squirt *Ciona intestinalis*, suggesting *eif4e1c* may have arisen in the tetrapod genome duplication that predates the teleost-specific event. Phylogenetic analysis was conducted using maximum likelihood as an optimality criterion as implemented in IQ-TREE2 (Minh et al., 2020). The JTT+G (4 state) model of amino acid substitution was selected using corrected Akaike Information Criterion (AIC) scores and used to estimate the phylogeny. Branch support was estimated using 1000 replicates of the ultrafast bootstrap with a minimum correlation coefficient of 0.99. IQ-TREE search parameters include a perturbation strength of 0.5 for the randomized nearest neighbor interchange with >100 unsuccessful iterations required to end the heuristic search of tree space. All analyses were conducted using the IQ-TREE web server (Trifinopoulos et al., 2016).

### Gene expression analysis from RNA sequencing

To examine *eif4e1c*, *eif4ea* and *eif4eb* expression in adult zebrafish tissues, we turned to previously published RNA sequencing datasets available on NCBI GEO. Breifly, fpkm was calculated from the raw read counts and normalized to the fpkm of *mob4* a gene that does not change expression between tissues (Hu et al., 2016). For zebrafish fins, we obtained data from 3 different studies (2 replicates from GSE126701 (Lee et al., 2020), 2 replicates from GSE146960: GSM 4411406 and GSM4411407 (Thompson et al., 2020), and 1 replicate from GSE76564 (Kang et al., 2016)). For zebrafish heart, we obtained datasets from 2 different studies (2 replicates from GSE75894 (Kang et al., 2016), 2 replicates from GSE168371: GSM5137453 and GSM5137454 (Pronobis et al., 2021)). Zebrafish brain, we looked at datasets from 2 different studies: 2 replicates from GSE158079- GSM4790340, GSM4790341(Sun et al., 2022) and 2 replicates from GSE151307 (Saraswathy et al., 2022). For zebrafish kidney, we obtained datasets from 2 different studies (2 replicates from GSE158079: GSM4790350 and GSM4790351, and 3 replicates from GSE193630: GSM5815254, GSM5815255, GSM5815256 (Sun et al., 2022)). The zebrafish liver datasets were derived from 1 study, GSE183023 (Heinkele et al., 2021) consisting of 6 replicates (GSM5549094). The spinal cord datasets were obtained from 2 studies: GSE183644 (2 replicates) (Saraswathy et al., 2022) and GSE77025 (1 replicate) (Mokalled et al., 2016). Only 2 replicates were available for testis, ovary, and oocyte from GSE111882 (Herberg et al., 2018).

### scRNA-seq analysis

For scRNA-seq analysis of uninjured and injured hearts, we obtained count files from GSE159032 and GSE158919 (Hu et al., 2020) and reanalyzed with Seurat package (Stuart et al., 2019). Low quality cells (nFeature_RNA ≥ 4100, mitoRatio ≥ 0.25) were filtered out. The 50 principal components (PCs) of the PCA with resolution 4 were used for clustering.

### Statistics

To ensure that experiments were sufficiently powered we used an α error probability of 0.05 and 80% power to calculate if sample sizes were sufficient. For each analysis we used the Welch’s parametric t-test to calculate significance. Outliers were removed using the ROUT (Robust regression and outlier removal) method available on the Graphpad PRISM software. To define outliers, PRISM uses the false discovery rate approach to handling multiple comparisons (Q = 1%) to remove the outliers, and then analyzes the data using ordinary least-squares regression. If the variance between the comparison groups was also significant (F test for unequal variances), we switched to using non-parametric test (Mann-Whitney). For the CM counting and CM proliferation assays, one researcher embedded hearts and sectioned slides and a separate researcher was blinded during imaging and quantification.

### Fish strains

All zebrafish (Danio rerio) used in this study derive from the Ekwill strain. Adults less than a year old were used for all the experiments with specific ages noted for the relevant experiments. Males and females were mixed in similar proportions in each of the conditions. All experiments were performed under university supervision according to the IACUC protocol #2018R00000090.

### Construction of *eif4e1c* deletion fish line

Guide RNAs (sgRNA) were designed against intron 1 (GGACATGTTTAAGCAGTGAG) and exon 7 (CAGAGGCACCTGCAGACTGA). DNA templates of the respective sgRNA fused to a tracRNA were produced by PCR. T7 transcribed sgRNA were then injected with Cas9 protein into newly fertilized embryos from EK parents. Adults injected at fertilization were mated and embryos of possible stables lines were screened by PCR oligos (Fwd: CGGTAAGAGGGATCGGGCTTGGT, Rev: GTCTACAAGCAAGATGCCACAATTCCAGCA) flanking the deleted region (chr13:18,311,769:18,323,899). The selected line, called *eif4e1c^pd370^* was a perfect deletion without addition of extra nucleotides between the fused cut sites. Heterozygote adults were mated as described. The Δ*eif4e1c* mutant line is available from the corresponding author upon request.

### Length and weight measurements

3-6mpf zebrafish were anesthetized in phenoxyethanol and their length was measured using a ruler starting from the eye to the center of caudal tail fin before the lobes diverge. After removing excess water with a Kimwipe, the anaesthetized zebrafish were weighed on an analytical scale. Fish were with different genotypes were raised and housed together and measured shortly after genotyping.

### Immuno-histochemistry

Primary antibodies used in this study were anti-Mef2 (1:100, Abcam ab197070), anti-hsEIF4E (1:100, Abcam - ab 33768), and anti-MF20 (1:100, DSHB MF20). Secondary antibodies were, anti-mouse Alexa546 (1:200) for MF20, and anti-rabbit-Alexa488 (1:200) for Mef2. Three sections representing the hearts with largest lumen were selected from each heart and imaged using a 20X objective on a Zeiss confocal. For obtaining cardiomyocyte counts, the number of Mef2c+ nuclei were counted using MIPAR image analysis software (REF Sosa). To calculate differences in immunofluorescence for anti-hsEIF4E, we drew regions of interest of the same size using ImageJ. The ratio of raw intensity (RawIntDen) was determined at the injury site versus an injury distal area after normalization to of background fluorescence. Phalloidin staining was performed according to the manufacturer’s protocol using 1:200 dilution (Abcam 176753). TUNEL assays were also performed according to manufacturer’s protocol (ThermoScientific C10617).

### Western blot

Originally raised against the human form of canonical EIF4E, we used an antibody (Abcam - ab33768) that targets a region highly homologous to both zebrafish canonical paralogs, Eif4ea and Eif4eb (95% similar and 82% identical, Fig. S6A). The regions in common are shared by all three proteins so the antibody should recognize both zebrafish paralogs. The antibody detects a major band of the expected size (∼25kD) by western when probed against whole cell extract from zebrafish hearts. In addition, the antibody demonstrated cytoplasmic staining patterns on zebrafish hearts that are consistent with what was found when used for immunofluorescence on human cells (Fig. S6B).

### OP-Puromycin experiments

OPP (Cayman Chemical) was resuspended in DMSO and then diluted with PBS to working solution of 5μM. Each fish was injected intraperitonially with 10μl and allowed to swim around for 1 hr before heart removal. Longer incubation times did not affect the data. Hearts were fixed, sectioned and then stained using CLICK chemistry with azide-594. To calculate fluorescence, images were opened in ImageJ and thresholding was used to calculate fluorescence. The average mean intensity was used for background readings to correct for total fluorescence. Three sections were averaged as technical replicates from each heart.

### Ribosome profiling

For each replicate sample, RNA from thirty zebrafish hearts was extracted for Ribo-seq library generation and parallel RNA sequencing. The hearts were lysed on ice with 100ul of lysis buffer with 53 mg/ml cycloheximide using mechanical homogenization with a pestle for thirty seconds. Cells were lysed further by trituration of the sample using a 27.5- gauge needle syringe 20x on ice. Cell extracts (average OD 7) were treated with MNase (NEB), CaCl_2_ (5mM final concentration), and Turbo DNAse I (Invitrogen) at 25°C for 30 minutes. MNase digestion was stopped with SUPERase-inhibitor incubation on ice. 80S ribosomes were isolated using SH400 spin columns (Cytiva). Columns were prepared according to the manufacturer’s instructions. 100 µls of MNase treated sample was loaded per column and centrifuged for 2 min at 600 RCF. The flow through was isolated and incubated for 5 min. with 3x volume of Trizol LS (Ambion). 200 µl of chloroform was added and incubated for 2 min. Samples were centrifuged for 15 min. at 12,000 rpm (4°C). The aqueous phase was extracted and precipitated overnight at −20°C using glycoblue (ThermoFisher) and isopropanol followed by centrifugation for 30 minutes at 4C at 10,000 RPM. Pellets were washed with 75% Ethanol and resuspended into RNase free water (Ingolia et al., 2012).

Samples were subjected to 15% denaturing polyacrylamide-mediated gel (UreaGel 29:1 gel system, National Diagnostics) and visualized by SYBR gold (ThermoFisher) after electrophoresis. Ribosome-protected RNA fragments 26-mers to 34-mers were excised and RNA was eluted overnight by rotating the gel slices in RNA extraction buffer (300 mM NaOAc (pH 5.2), 1mM EDTA, 0.25% (w/v) SDS at room temperature. To deplete rRNA fragments, the isolated RNA was precipitated on dry ice for 30 minutes using glycoblue (ThermoFisher) and isopropanol and treated with the RiboMinus Ribosomal RNA depletion kit (Ambion), according to the manufacturer’s instructions.

The RNA samples were then de-phosphorylated by adding 1ul of T4 PNK, 1ul SUPERase-In, 7ul of 10X PNK reaction buffer for a 70ul volume reaction. The reaction is incubated then for 1 hour at 37C before adding 1ul of 100mM ATP for 30 minutes at 37C. The reaction is then heat inactivated at 70C for 10 minutes and precipitated as before. Adapter ligation, reverse transcription and subsequent cDNA amplification were performed using the NEXT-Flex smRNA-Seq Kit v3 (Perkin Elmer). Samples were subjected to 8% native PAGE (0.5X TBE) gel electrophoresis. cDNA products of the final PCR were visualized by SYBR gold (ThermoFisher), excised from the gels, and extracted overnight with DNA extraction buffer (300 mM NaCl, 10mM Tris (pH 8.0), 1 mM EDTA) and precipitated as described previously.

To prepare RNA sequencing libraries, total RNA was isolated from each sample using Trizol LS as described above and subjected to Turbo DNase I digestion (1ul Turbo Dnase 1, 1x DNase reaction buffer, 20 minutes at 37C). RNA sequencing libraries were constructed from 500ng of purified RNA using Illumina’s TruSeq Stranded Total RNA kit, according to the manufacturer’s instructions, by the Case Western Reserve University (CWRU) Genomics Core.

Libraries were assessed for quality by Agilent Bioanlyzer followed subsequently by single-end 75 cycle sequencing. The RNA sequencing and Ribo-seq sequencing were performed at the CWRU Genomics Core on the Illumina NEXTseq 550 v2.5 (High Output) platform. Raw and processed data are available at GEO (GSE Accession #211793).

### Bioinformatics

Ribosome profiling – Fastq files were processed using FastQC (v0.11.8; RRID:SCR_014583) for quality control (QC) analysis. Pre-processing of sequencing reads was based on the QC report using the FASTQ Quality Filter module in the FASTX-Toolkit (RRID:SCR_019035). This was used to filter the bases with 99% accuracy based on Q Score. Reads were removed that showed less than 70% of nucleotides with an accuracy of at least 99%. The sequences were collapsed according to their Unique Molecular Identifiers (UMIs). Cutadapt (RRID:SCR_011841) was used to trim 21-nt adaptor before the first nt and the last 4nt from the reads to remove the UMI.

The processed reads were aligned to GRCz11/danRer11 (assembly of the zebrafish genome; Genome Reference Consortium) using STAR v2.5.3a (RRID:SCR_004463). Sequencing replicates were highly similar to one another for both ribosome protected fragments and input mRNA (Fig. S5A). Generated bam files were processed with RiboProfiling package v1.2.2 (Popa et al., 2016). Coverage counts on the coding regions (CDS) were obtained for each sample based on RiboProfiling function modules, *TxDb.Drerio.UCSC.danRer11.refGene (*v3.4.6; Annotation package for TxDb object(s); Team BC, 2019) and GenomicFeatures package v1.46.1 (Lawrence et al., 2013). Differential expression analysis was performed using the DESeq2 pipeline (version 1.26.0), based on the expressed raw reads (Love et al., 2014). Statistic thresholds for the features in the comparison results (p-values and log2 fold changes), were used for the gene extraction, as described.

Genes with transcripts with increasing translation efficiency in mutant hearts were calculated as (1) significantly increasing by RPF and not mRNA, (2) not significantly increasing by RPF but significantly decreasing by mRNA or (3) significantly changing in both datasets with an RPF/mRNA ratio > 1. Genes with transcripts translated less efficiently were identified as (1) decreasing significantly by RPF and not changing by mRNA, (2) not significantly changing in RPF but significantly increasing in mRNA, or (3) significantly changing in both datasets with an RPF/mRNA ratio < 1. Gene ontology categories were identified using DAVID (Huang et al., 2009; Sherman et al., 2022).

### Zebrafish cardiac regeneration experiments

Zebrafish were anesthetized using Tricaine and placed ventral side up on a sponge to carry out resection of the ventricular apex. Iridectomy scissors were used to make an incision through the skin and pericardial sac. Gentle abdominal pressure exposed the heart and ∼20% of the apex was removed with scissors, penetrating the chamber lumen (Poss et al., 2002). Hearts were harvested 7, 14 or 30 days after injury depending on the experiment. To genetically ablate cardiomyocytes, *cmlc2:CreER^pd10^; bactin2:loxp-mCherry-STOP-loxp-DTA^pd36^* (Z-CAT) fish were incubated in 0.5 μM tamoxifen for 17 hours (Wang et al., 2011).

#### To quantify CM proliferation

Injured fish were injected into the abdominal cavity once every 24hrs for 3days (4-6dpa) with 10ul of a 10mM solution of EdU diluted in PBS. Hearts were removed on day 7, embedded and cryo-sectioned. Slides were stained with Alexa594-azide using CLICK chemistry and then immunostained for Mef2c. Briefly, sections were blocked with 1% BSA (Fraction V) and 5% goat serum and washed in PBS with .2% triton-100. Three sections representing the largest wound area were selected from each heart and imaged using a 20X objective. The number of Mef2+ and Mef2+EdU+ cells were counted using the MIPAR image analysis software and the CM proliferation index was calculated as the number of Mef2+EdU+ cells / total Mef2+ cells (Sosa et al., 2014). The CM proliferation index was averaged across 2- 4 appropriate sections from each heart.

#### To visualize regrowth

Hearts were removed 28dpa, embedded and cryo-sectioned. Slides were incubated at 60 °C for 3 h in Bouin-fixative previously heated at 60 °C for 30 min. Slides were then rinsed in distilled water (dH_2_O) for 30 min, and then incubated with 1% (w/v) phosphomolybdic acid solution for 5 min. Following a 5 min wash-step in dH_2_O, sections were stained with the AFOG-solution for 5 min, washed, dehydrated, and then mounted in Cytoseal. Three sections representing the largest wound area were selected from each heart and imaged using a 20X objective.

### Fin regeneration analysis

Fish were anesthetized in phenoxyethanol, and the caudal fins were amputated just below bifurcations of rays using a razor blade and removing ∼1/2 of the fin from body to tip. Animals were allowed to regenerate for 4 days after which fins were imaged using a 10X objective. Fin regrowth was measured from the amputation till the tip of each ray using the Zeiss microscope software. The 2^nd^ through 4th rays were averaged from both the dorsal and ventral sides.

### Seahorse analysis of mitochondrial respiration

Mitochondria from fish hearts were isolated as previously described in mice (Peri-Okonny et al., 2019). Oxygen consumption rates were optimized for 1.5μg of mitochondria which were resuspended in respiratory buffer (70mM sucrose, 220mM mannitol, 10mM K2HPO4, 5mM MgCl2, 2mM HEPES, 1mM EGTA, pH 7.4. Electron coupling (EC) assays were performed using the Seahorse Bioanalyzer (Agilent). The EC assay measures basal respiration in coupled mitochondria with substrates present (succinate/rotenone), but no ADP, which drives respiration through electron transport chain (ETC) complexes II-IV. Active respiration is initiated by addition of ADP to mitochondria, increasing OCR. Oligomycin, an ATP synthase/Complex V inhibitor, decreases OCR. FCCP, an ETC chemical uncoupler, allows for maximal uncoupled respiration. Antimycin A, which inhibits complex III, shut down electron flow, ATP production, and OCR. Proton leak through the ETC can be determined with oligomycin treatment, and spare respiratory capacity is the difference in maximal respiration (FCCP treatment) and protein leak (Oligomycin treatment). After Seahorse measurements, the abundance of mitochondria was verified using qPCR and used to normalize the readings.

### SDH staining

Hearts were embedded fresh and sectioned and stained the same day. The succinate dehydrogenase (Sdh) enzymatic activity was measured by incubating sections with 37.5 mM sodium phosphate buffer pH 7.60, 70 mM sodium succinate, 5 mM sodium azide and 0.4 mM tetranitro blue tetrazolium (TNBT) for 20 min at 28°C (Honkoop et al., 2018). The reaction was quenched in 10 mM HCl, and slides were mounted in glycerin-gelatin mounting media. Images were processed with ImageJ calculating percent mean intensity of signal averaged by the total area of the heart (pixels).

### Gene expression analysis by RT-qPCR

Dissected tissues were placed in Hank’s Buffered Salt Solution and washed three times in PBS before being snap frozen. For regenerating fins, only the blastema was removed. Organs from 10 fish were pooled (5 male, 5 female). Primer pair efficiency was assessed and validated to be 95%-105%. RNA was isolated using an adapted version of a published protocol and then DNased. Reverse transcription (RT) was performed with 1ug total RNA using SuperscriptIII (Thermo) and qPCR was performed with 3 technical replicates for each biological replicate. Controls without RNA template in the RT were run for each primer pair to identify possible DNA contamination and none was found. Fold gene expression was determined using the ΔΔCt method and normalized to *mob4* (Hu et al., 2016). At least three biological replicates were used for each condition.

## Acknowledgements

We would like to thank the Busch-Nentwich lab for providing RNA-seq data from zebrafish embryos (http://www.ebi.ac.uk/gxa/experiments/E-ERAD-475). Confocal imaging was performed in the Neuroscience Imaging Core at OSU, made possible by NIH Shared Instrumentation Grant (S10 OD026842). Thanks to Charles Bell and Tianmin Fu who helped guide the structural modeling. We acknowledge funding from NIH grant R35HL150713 for K.D.P., NIH grants R35GM137878 and R01HL151522 for J.K., and from an American Heart Association grant 17SDG33660922 and the Department of Biological Chemistry and Pharmacology for J.A.G.

## REFERENCES

Altmann, M., Müller, P. P., Pelletier, J., Sonenberg, N. and Trachsel, H. (1989). A Mammalian Translation Initiation Factor Can Substitute for Its Yeast Homologue in Vivo. J Biol Chem 264, 12145–12147.

Austin, C. M., Tan, M. H., Croft, L. J., Hammer, M. P. and Gan, H. M. (2015). Whole Genome Sequencing of the Asian Arowana (Scleropages formosus) Provides Insights into the Evolution of Ray-Finned Fishes. Genome Biol Evol 7, 2885–2895.

Baek, M., DiMaio, F., Anishchenko, I., Dauparas, J., Ovchinnikov, S., Lee, G. R., Wang, J., Cong, Q., Kinch, L. N., Schaeffer, R. D., et al. (2021). Accurate prediction of protein structures and interactions using a 3-track neural network. Sci New York N Y 373, 871–876.

Bezawork-Geleta, A., Rohlena, J., Dong, L., Pacak, K. and Neuzil, J. (2017). Mitochondrial Complex II: At the Crossroads. Trends Biochem Sci 42, 312–325.

Bian, C., Hu, Y., Ravi, V., Kuznetsova, I. S., Shen, X., Mu, X., Sun, Y., You, X., Li, J., Li, X., et al. (2016). The Asian arowana (Scleropages formosus) genome provides new insights into the evolution of an early lineage of teleosts. Sci Rep-uk 6, 24501.

Borden, K. L. B. and Volpon, L. (2020). The diversity, plasticity, and adaptability of cap-dependent translation initiation and the associated machinery. Rna Biol 17, 1–13.

Braasch, I., Gehrke, A. R., Smith, J. J., Kawasaki, K., Manousaki, T., Pasquier, J., Amores, A., Desvignes, T., Batzel, P., Catchen, J., et al. (2016). The spotted gar genome illuminates vertebrate evolution and facilitates human-teleost comparisons. Nat Genet 48, 427 437.

Buccitelli, C. and Selbach, M. (2020). mRNAs, proteins and the emerging principles of gene expression control. Nat Rev Genet 21, 630–644.

Cardoso, A. C., Lam, N. T., Savla, J. J., Nakada, Y., Pereira, A. H. M., Elnwasany, A., Menendez-Montes, I., Ensley, E. L., Petric, U. B., Sharma, G., et al. (2020). Mitochondrial Substrate Utilization Regulates Cardiomyocyte Cell Cycle Progression. Nat Metabolism 2, 167–178.

Chen, B.-R., Wei, T.-W., Tang, C.-P., Sun, J.-T., Shan, T.-K., Fan, Y., Yang, T.-T., Li, Y.-F., Ma, Y., Wang, S.-B., et al. (2022). MNK2-eIF4E axis promotes cardiac repair in the infarcted mouse heart by activating cyclin D1. J Mol Cell Cardiol 166, 91–106.

Chorghade, S., Seimetz, J., Emmons, R., Yang, J., Bresson, S. M., Lisio, M. D., Parise, G., Conrad, N. K. and Kalsotra, A. (2017). Poly(A) tail length regulates PABPC1 expression to tune translation in the heart. Elife 6, 568.

Davis, M. R., Delaleau, M. and Borden, K. L. B. (2019). Nuclear eIF4E Stimulates 3’-End Cleavage of Target RNAs. Cell Reports 27, 1397 1408.e4.

Echtay, K. S., Roussel, D., St-Pierre, J., Jekabsons, M. B., Cadenas, S., Stuart, J. A., Harper, J. A., Roebuck, S. J., Morrison, A., Pickering, S., et al. (2002). Superoxide activates mitochondrial uncoupling proteins. Nature 415, 96–99.

Emmott, E., Jovanovic, M. and Slavov, N. (2018). Ribosome Stoichiometry: From Form to Function. Trends Biochem Sci 44, 95–109.

Fang, Y., Gupta, V., Karra, R., Holdway, J. E., Kikuchi, K. and Poss, K. D. (2013). Translational profiling of cardiomyocytes identifies an early Jak1/Stat3 injury response required for zebrafish heart regeneration. Proc National Acad Sci 110, 13416 13421.

Fukuda, R., Marín-Juez, R., El-Sammak, H., Beisaw, A., Ramadass, R., Kuenne, C., Guenther, S., Konzer, A., Bhagwat, A. M., Graumann, J., et al. (2020). Stimulation of glycolysis promotes cardiomyocyte proliferation after injury in adult zebrafish. Embo Rep 21, e49752.

Genuth, N. R. and Barna, M. (2018). Heterogeneity and specialized functions of translation machinery: from genes to organisms. Nat Rev Genet 19, 431 452.

Goldman, J. A., Kuzu, G., Lee, N., Karasik, J., Gemberling, M., Foglia, M. J., Karra, R., Dickson, A. L., Sun, F., Tolstorukov, M. Y., et al. (2017). Resolving Heart Regeneration by Replacement Histone Profiling. Dev Cell 40, 392 404.e5.

Graff, J. R., Konicek, B. W., Vincent, T. M., Lynch, R. L., Monteith, D., Weir, S. N., Schwier, P., Capen, A., Goode, R. L., Dowless, M. S., et al. (2007). Therapeutic suppression of translation initiation factor eIF4E expression reduces tumor growth without toxicity. J Clin Invest 117, 2638– 2648.

Grüner, S., Peter, D., Weber, R., Wohlbold, L., Chung, M.-Y., Weichenrieder, O., Valkov, E., Igreja, C. and Izaurralde, E. (2016). The Structures of eIF4E-eIF4G Complexes Reveal an Extended Interface to Regulate Translation Initiation. Mol Cell 64, 467–479.

Harvey, S. A., Sealy, I., Kettleborough, R., Fenyes, F., White, R., Stemple, D. and Smith, J. C. (2013). Identification of the zebrafish maternal and paternal transcriptomes. Dev Camb Engl 140, 2703–2710.

Heinkele, F. J., Lou, B., Erben, V., Bennewitz, K., Poschet, G., Sticht, C. and Kroll, J. (2021). Metabolic and Transcriptional Adaptations Improve Physical Performance of Zebrafish. Antioxidants Basel Switz 10, 1581.

Herberg, S., Gert, K. R., Schleiffer, A. and Pauli, A. (2018). The Ly6/uPAR protein Bouncer is necessary and sufficient for species-specific fertilization. Science 361, 1029–1033.

Hirose, K., Payumo, A. Y., Cutie, S., Hoang, A., Zhang, H., Guyot, R., Lunn, D., Bigley, R. B., Yu, H., Wang, J., et al. (2019). Evidence for hormonal control of heart regenerative capacity during endothermy acquisition. Science 364, 184 188.

Honkoop, H., Bakker, D. E. de, Aharonov, A., Kruse, F., Shakked, A., Nguyen, P. D., Heus, C. de, Garric, L., Muraro, M. J., Shoffner, A., et al. (2018). Single-cell analysis uncovers that metabolic reprogramming by ErbB2 signaling is essential for cardiomyocyte proliferation in the regenerating heart. Elife 8, e50163.

Hu, Y., Xie, S. and Yao, J. (2016). Identification of Novel Reference Genes Suitable for qRT-PCR Normalization with Respect to the Zebrafish Developmental Stage. Plos One 11, e0149277.

Hu, B., Lelek, S., Spanjaard, B., El-Sammak, H., Simões, M. G., Mintcheva, J., Aliee, H., Schäfer, R., Meyer, A. M., Theis, F., et al. (2022). Origin and function of activated fibroblast states during zebrafish heart regeneration. Nat Genet 54, 1227–1237.

Huang, D. W., Sherman, B. T. and Lempicki, R. A. (2009). Systematic and integrative analysis of large gene lists using DAVID bioinformatics resources. Nat Protoc 4, 44–57.

Ingolia, N. T., Brar, G. A., Rouskin, S., McGeachy, A. M. and Weissman, J. S. (2012). The ribosome profiling strategy for monitoring translation in vivo by deep sequencing of ribosome-protected mRNA fragments. Nat Protoc 7, 1534–1550.

Joshi, B., Lee, K., Maeder, D. L. and Jagus, R. (2005). Phylogenetic analysis of eIF4E-family members. Bmc Evol Biol 5, 48–48.

Jumper, J., Evans, R., Pritzel, A., Green, T., Figurnov, M., Ronneberger, O., Tunyasuvunakool, K., Bates, R., Žídek, A., Potapenko, A., et al. (2021). Highly accurate protein structure prediction with AlphaFold. Nature 596, 583–589.

Kang, J., Hu, J., Karra, R., Dickson, A. L., Tornini, V. A., Nachtrab, G., Gemberling, M., Goldman, J. A., Black, B. L. and Poss, K. D. (2016). Modulation of tissue repair by regeneration enhancer elements. Nature 532, 201 206.

Keegan, B. R., Feldman, J. L., Begemann, G., Ingham, P. W. and Yelon, D. (2005). Retinoic acid signaling restricts the cardiac progenitor pool. Science 307, 247 249.

Kikuchi, K., Holdway, J. E., Major, R. J., Blum, N., Dahn, R. D., Begemann, G. and Poss, K. D. (2011). Retinoic acid production by endocardium and epicardium is an injury response essential for zebrafish heart regeneration. Dev Cell 20, 397 404.

Kong, J. and Lasko, P. (2012). Translational control in cellular and developmental processes. Nat Rev Genet 13, 383–394.

Kubacka, D., Miguel, R. N., Minshall, N., Darzynkiewicz, E., Standart, N. and Zuberek, J. (2015). Distinct Features of Cap Binding by eIF4E1b Proteins. J Mol Biol 427, 387–405.

Kusnadi, E. P., Timpone, C., Topisirovic, I., Larsson, O. and Furic, L. (2021). Regulation of gene expression via translational buffering. Biochimica Et Biophysica Acta Bba - Mol Cell Res 1869, 119140.

Lai, S.-L., Marín-Juez, R., Moura, P. L., Kuenne, C., Lai, J. K. H., Tsedeke, A. T., Guenther, S., Looso, M. and Stainier, D.Y. (2017). Reciprocal analyses in zebrafish and medaka reveal that harnessing the immune response promotes cardiac regeneration. Elife 6, 1382.

Lawrence, M., Huber, W., Pagès, H., Aboyoun, P., Carlson, M., Gentleman, R., Morgan, M. T. and Carey, V. J. (2013). Software for Computing and Annotating Genomic Ranges. Plos Comput Biol 9, e1003118.

Lee, H. J., Hou, Y., Chen, Y., Dailey, Z. Z., Riddihough, A., Jang, H. S., Wang, T. and Johnson, S. L. (2020). Regenerating zebrafish fin epigenome is characterized by stable lineage-specific DNA methylation and dynamic chromatin accessibility. Genome Biol 21, 52.

Lepilina, A., Coon, A. N., Kikuchi, K., Holdway, J. E., Roberts, R. W., Burns, C. G. and Poss, K. D. (2006). A dynamic epicardial injury response supports progenitor cell activity during zebrafish heart regeneration. Cell 127, 607 619.

Lin, J.-J., Wang, F.-Y., Li, W.-H. and Wang, T.-Y. (2017). The rises and falls of opsin genes in 59 ray-finned fish genomes and their implications for environmental adaptation. Sci Rep-uk 7, 15568.

Lorent, J., Kusnadi, E. P., Hoef, V., Rebello, R. J., Leibovitch, M., Ristau, J., Chen, S., Lawrence, M. G., Szkop, K. J., Samreen, B., et al. (2019). Translational offsetting as a mode of estrogen receptor α-dependent regulation of gene expression. Embo J 38, e101323.

Love, M. I., Huber, W. and Anders, S. (2014). Moderated estimation of fold change and dispersion for RNA-seq data with DESeq2. Genome Biol 15, 550.

Ma, D., Tu, C., Sheng, Q., Yang, Y., Kan, Z., Guo, Y., Shyr, Y., Scott, I. C. and Lou, X. (2018). Dynamics of Zebrafish Heart Regeneration Using an HPLC–ESI–MS/MS Approach. J Proteome Res 17, 1300 1308.

Miklas, J. W., Levy, S., Hofsteen, P., Mex, D. I., Clark, E., Muster, J., Robitaille, A. M., Sivaram, G., Abell, L., Goodson, J. M., et al. (2021). Amino acid primed mTOR activity is essential for heart regeneration. Iscience 25, 103574.

Minh, B. Q., Schmidt, H. A., Chernomor, O., Schrempf, D., Woodhams, M. D., Haeseler, A. von and Lanfear, R. (2020). IQ-TREE 2: New Models and Efficient Methods for Phylogenetic Inference in the Genomic Era. Mol Biol Evol 37, 1530–1534.

Minshall, N., Reiter, M. H., Weil, D. and Standart, N. (2007). CPEB Interacts with an Ovary-specific eIF4E and 4E-T in Early Xenopus Oocytes*. J Biol Chem 282, 37389–37401.

Mokalled, M. H., Patra, C., Dickson, A. L., Endo, T., Stainier, D. Y. R. and Poss, K. D. (2016). Injury-induced ctgfa directs glial bridging and spinal cord regeneration in zebrafish. Science 354, 630 634.

Okuda, A. and Kimura, G. (1996). An Amino Acid Change in Novel Protein D123 Is Responsible for Temperature-Sensitive G1-Phase Arrest in a Mutant of Rat Fibroblast Line 3Y1. Exp Cell Res 223, 242–249.

Peri-Okonny, P. A., Baskin, K. K., Iwamoto, G., Mitchell, J. H., Smith, S. A., Kim, H. K., Szweda, L. I., Bassel-Duby, R., Fujikawa, T., Castorena, C. M., et al. (2019). High-Phosphate Diet Induces Exercise Intolerance and Impairs Fatty Acid Metabolism in Mice. Circulation 139, 1422–1434.

Perzlmaier, A. F., Richter, F. and Seufert, W. (2013). Translation Initiation Requires Cell Division Cycle 123 (Cdc123) to Facilitate Biogenesis of the Eukaryotic Initiation Factor 2 (eIF2). J Biol Chem 288, 21537–21546.

Popa, A., Lebrigand, K., Paquet, A., Nottet, N., Robbe-Sermesant, K., Waldmann, R. and Barbry, P. (2016). RiboProfiling: a Bioconductor package for standard Ribo-seq pipeline processing. F1000research 5, 1309.

Poss, K. D., Wilson, L. G. and Keating, M. T. (2002). Heart regeneration in zebrafish. Science 298, 2188 2190.

Pronobis, M. I., Zheng, S., Singh, S. P., Goldman, J. A. and Poss, K. D. (2021). In vivo proximity labeling identifies cardiomyocyte protein networks during zebrafish heart regeneration. Elife 10, e66079.

Robalino, J., Joshi, B., Fahrenkrug, S. C. and Jagus, R. (2004). Two Zebrafish eIF4E Family Members Are Differentially Expressed and Functionally Divergent*. J Biol Chem 279, 10532–10541.

Sakurai, T., Tsuchida, M., Lampe, P. D. and Murakami, M. (2013). Cardiomyocyte FGF signaling is required for Cx43 phosphorylation and cardiac gap junction maintenance. Exp Cell Res 319, 2152– 2165.

Saraswathy, V. M., Zhou, L., McAdow, A. R., Burris, B., Dogra, D., Reischauer, S. and Mokalled, M.H. (2022). Myostatin is a negative regulator of adult neurogenesis after spinal cord injury in zebrafish. Cell Reports 41, 111705.

Sénéchal, P., Robert, F., Cencic, R., Yanagiya, A., Chu, J., Sonenberg, N., Paquet, M. and Pelletier, J. (2021). Assessing eukaryotic initiation factor 4F subunit essentiality by CRISPR-induced gene ablation in the mouse. Cell Mol Life Sci 78, 6709–6719.

Sherman, B. T., Hao, M., Qiu, J., Jiao, X., Baseler, M. W., Lane, H. C., Imamichi, T. and Chang, W. (2022). DAVID: a web server for functional enrichment analysis and functional annotation of gene lists (2021 update). Nucleic Acids Res 50, W216–W221.

Sosa, J. M., Huber, D. E., Welk, B. and Fraser, H. L. (2014). Development and application of MIPAR^TM^: a novel software package for two-and three-dimensional microstructural characterization. Integrating Mater Manuf Innovation 3, 123–140.

Stockdale, W. T., Lemieux, M. E., Killen, A. C., Zhao, J., Hu, Z., Riepsaame, J., Hamilton, N., Kudoh, T., Riley, P. R., Aerle, R. van, et al. (2018). Heart Regeneration in the Mexican Cavefish. Cell Reports 25, 1997 2007.e7.

Stuart, T., Butler, A., Hoffman, P., Hafemeister, C., Papalexi, E., Mauck, W. M., Hao, Y., Stoeckius, M., Smibert, P. and Satija, R. (2019). Comprehensive Integration of Single-Cell Data. Cell 177, 1888–1902.e21.

Sun, L., Gao, J., Dong, X., Liu, M., Li, D., Shi, X., Dong, J.-T., Lu, X., Liu, C. and Zhou, J. (2008). EB1 promotes Aurora-B kinase activity through blocking its inactivation by protein phosphatase 2A. Proc National Acad Sci 105, 7153–7158.

Sun, F., Ou, J., Shoffner, A. R., Luan, Y., Yang, H., Song, L., Safi, A., Cao, J., Yue, F., Crawford, G. E., et al. (2022). Enhancer selection dictates gene expression responses in remote organs during tissue regeneration. Nat Cell Biol 24, 685–696.

Syntichaki, P., Troulinaki, K. and Tavernarakis, N. (2007). eIF4E function in somatic cells modulates ageing in Caenorhabditis elegans. Nature 445, 922–926.

Tahara, N., Akiyama, R., Wang, J., Kawakami, H., Bessho, Y. and Kawakami, Y. (2021). The FGF- AKT pathway is necessary for cardiomyocyte survival for heart regeneration in zebrafish. Dev Biol 472, 30–37.

Taylor, J. S., Peer, Y. V. de, Braasch, I. and Meyer, A. (2001). Comparative genomics provides evidence for an ancient genome duplication event in fish. Philosophical Transactions Royal Soc Lond Ser B Biological Sci 356, 1661–1679.

Thompson, J. D., Ou, J., Lee, N., Shin, K., Cigliola, V., Song, L., Crawford, G. E., Kang, J. and Poss, K. D. (2020). Identification and requirements of enhancers that direct gene expression during zebrafish fin regeneration. Development 147, dev191262.

Trifinopoulos, J., Nguyen, L.-T., Haeseler, A. von and Minh, B. Q. (2016). W-IQ-TREE: a fast online phylogenetic tool for maximum likelihood analysis. Nucleic Acids Res 44, W232–W235.

Truitt, M. L., Conn, C. S., Shi, Z., Pang, X., Tokuyasu, T., Coady, A. M., Seo, Y., Barna, M. and Ruggero, D. (2015). Differential Requirements for eIF4E Dose in Normal Development and Cancer. Cell 162, 59 71.

Wang, J., Panáková, D., Kikuchi, K., Holdway, J. E., Gemberling, M., Burris, J. S., Singh, S. P., Dickson, A. L., Lin, Y.-F., Sabeh, M. K., et al. (2011). The regenerative capacity of zebrafish reverses cardiac failure caused by genetic cardiomyocyte depletion. Development 138, 3421 3430.

Wang, Z.-Y., Leushkin, E., Liechti, A., Ovchinnikova, S., Mößinger, K., Brüning, T., Rummel, C., Grützner, F., Cardoso-Moreira, M., Janich, P., et al. (2020a). Transcriptome and translatome co-evolution in mammals. Nature 588, 642–647.

Wang, W., Hu, C.-K., Zeng, A., Alegre, D., Hu, D., Gotting, K., Granillo, A. O., Wang, Y., Robb, S., Schnittker, R., et al. (2020b). Changes in regeneration-responsive enhancers shape regenerative capacities in vertebrates. Science 369, eaaz3090.

White, R. J., Collins, J. E., Sealy, I. M., Wali, N., Dooley, C. M., Digby, Z., Stemple, D. L., Murphy, D. N., Billis, K., Hourlier, T., et al. (2017). A high-resolution mRNA expression time course of embryonic development in zebrafish. Elife 6, e30860.

Wu, C.-C., Kruse, F., Vasudevarao, M. D., Junker, J. P., Zebrowski, D. C., Fischer, K., Noël, E. S., Grün, D., Berezikov, E., Engel, F. B., et al. (2016). Spatially Resolved Genome-wide Transcriptional Profiling Identifies BMP Signaling as Essential Regulator of Zebrafish Cardiomyocyte Regeneration. Dev Cell 36, 36–49.

You, X., Bian, C., Zan, Q., Xu, X., Liu, X., Chen, J., Wang, J., Qiu, Y., Li, W., Zhang, X., et al. (2014). Mudskipper genomes provide insights into the terrestrial adaptation of amphibious fishes. Nat Commun 5, 5594.

Zhang, Y., Qin, C., Yang, L., Lu, R., Zhao, X. and Nie, G. (2018). A comparative genomics study of carbohydrate/glucose metabolic genes: from fish to mammals. Bmc Genomics 19, 246.

