## Supplemental Figures for "The Translation Initiation Factor Homolog, *eif4e1c*, Regulates Cardiomyocyte Metabolism and Proliferation During Heart Regeneration"

### Figure S1A

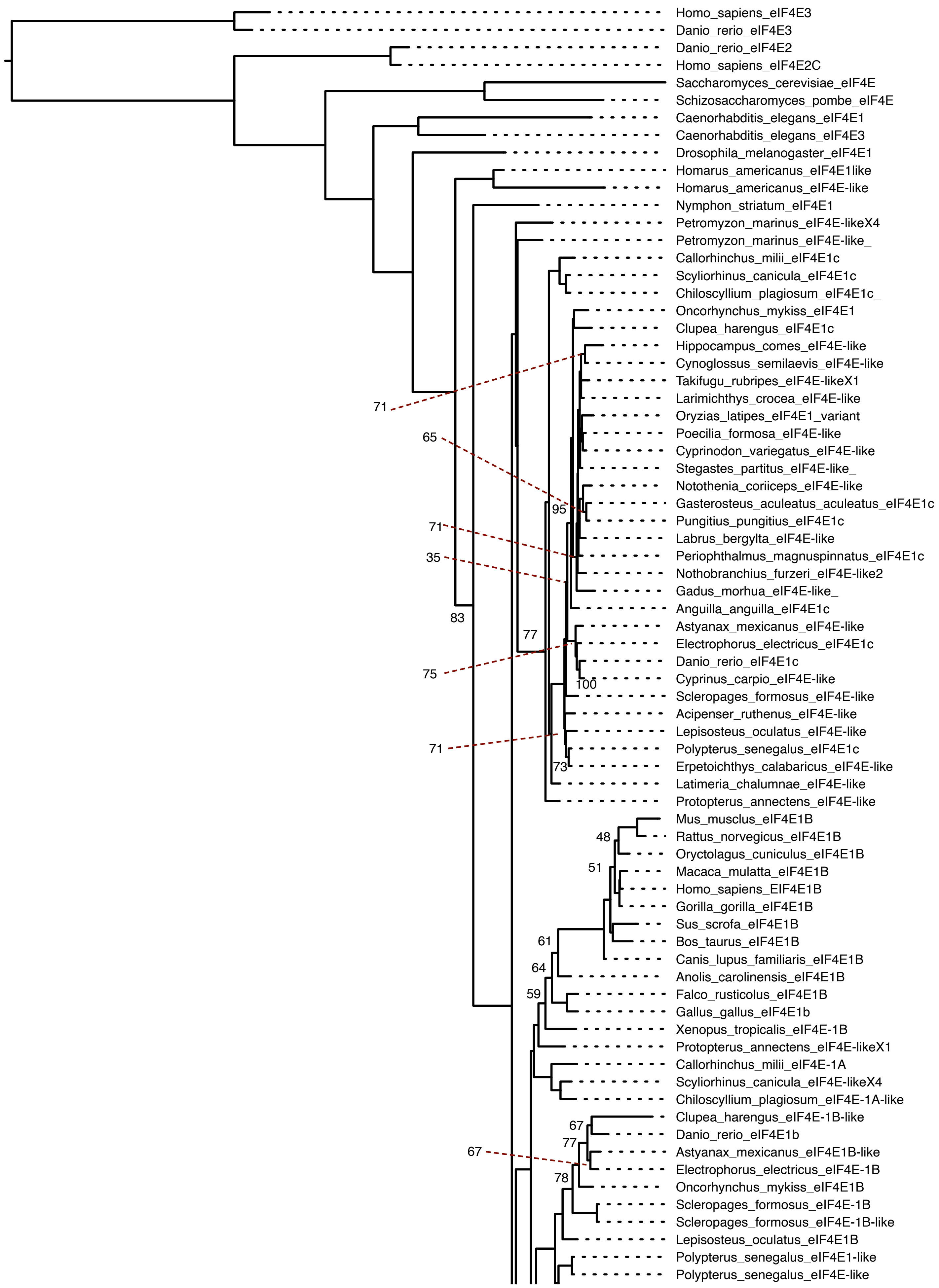

Homo_sapiens_eIF4E3
Danio_rerio_eIF4E3
Danio_rerio_eIF4E2
Homo_sapiens_eIF4E2C
Saccharomyces_cerevisiae_eIF4E
Schizosaccharomyces_pombe_eIF4E
Caenorhabditis_elegans_eIF4E1
Caenorhabditis_elegans_eIF4E3
Drosophila_melanogaster_eIF4E1
Homarus_americanus_eIF4E1like
Homarus_americanus_eIF4E-like
Nymphon_striatum_eIF4E1
Petromyzon_marinus_eIF4E-likeX4
Petromyzon_marinus_eIF4E-like_
Callorhinchus_milii_eIF4E1c
Scyliorhinus_canicula_eIF4E1c
Chiloscyllium_plagiosum_eIF4E1c_
Oncorhynchus_mykiss_eIF4E1
Clupea_harengus_eIF4E1c
Hippocampus_comes_eIF4E-like
Cynoglossus_semilaevis_eIF4E-like
Takifugu_rubripes_eIF4E-likeX1
Larimichthys_crocea_eIF4E-like
Oryzias_latipes_eIF4E1_variant
Poecilia_formosa_eIF4E-like
Cyprinodon_variegatus_eIF4E-like
Stegastes_partitus_eIF4E-like_
Notothenia_coriiceps_eIF4E-like
Gasterosteus_aculeatus_aculeatus_eIF4E1c
Pungitius_pungitius_eIF4E1c
Labrus_bergylta_eIF4E-like
Periophthalmus_magnuspinnatus_eIF4E1c
Nothobranchius_furzeri_eIF4E-like2
Gadus_morhua_eIF4E-like_
Anguilla_anguilla_eIF4E1c
Astyanax_mexicanus_eIF4E-like
Electrophorus_electricus_eIF4E1c
Danio_rerio_eIF4E1c
Cyprinus_carpio_eIF4E-like
Scleropages_formosus_eIF4E-like
Acipenser_ruthenus_eIF4E-like
Lepisosteus_oculatus_eIF4E-like
Polypterus_senegalus_eIF4E1c
Erpetoichthys_calabaricus_eIF4E-like
Latimeria_chalumnae_eIF4E-like
Protopterus_annectens_eIF4E-like
Mus_musclus_eIF4E1B
Rattus_norvegicus_eIF4E1B
Oryctolagus_cuniculus_eIF4E1B
Macaca_mulatta_eIF4E1B
Homo_sapiens_EIF4E1B
Gorilla_gorilla_eIF4E1B
Sus_scrofa_eIF4E1B
Bos_taurus_eIF4E1B
Canis_lupus_familiaris_eIF4E1B
Anolis_carolinensis_eIF4E1B
Falco_rusticolus_eIF4E1B
Gallus_gallus_eIF4E1b
Xenopus_tropicalis_eIF4E-1B
Protopterus_annectens_eIF4E-likeX1
Callorhinchus_milii_eIF4E-1A
Scyliorhinus_canicula_eIF4E-likeX4
Chiloscyllium_plagiosum_eIF4E-1A-like
Clupea_harengus_eIF4E-1B-like
Danio_rerio_eIF4E1b
Astyanax_mexicanus_eIF4E1B-like
Electrophorus_electricus_eIF4E-1B
Oncorhynchus_mykiss_eIF4E1B
Scleropages_formosus_eIF4E-1B
Scleropages_formosus_eIF4E-1B-like
Lepisosteus_oculatus_eIF4E1B
Polypterus_senegalus_eIF4E1-like
Polypterus_senegalus_eIF4E-like
71
65
95
71
35
83
77
75
100
71
73
48
51
61
64
59
67
77
67
78

### Figure S1A continued

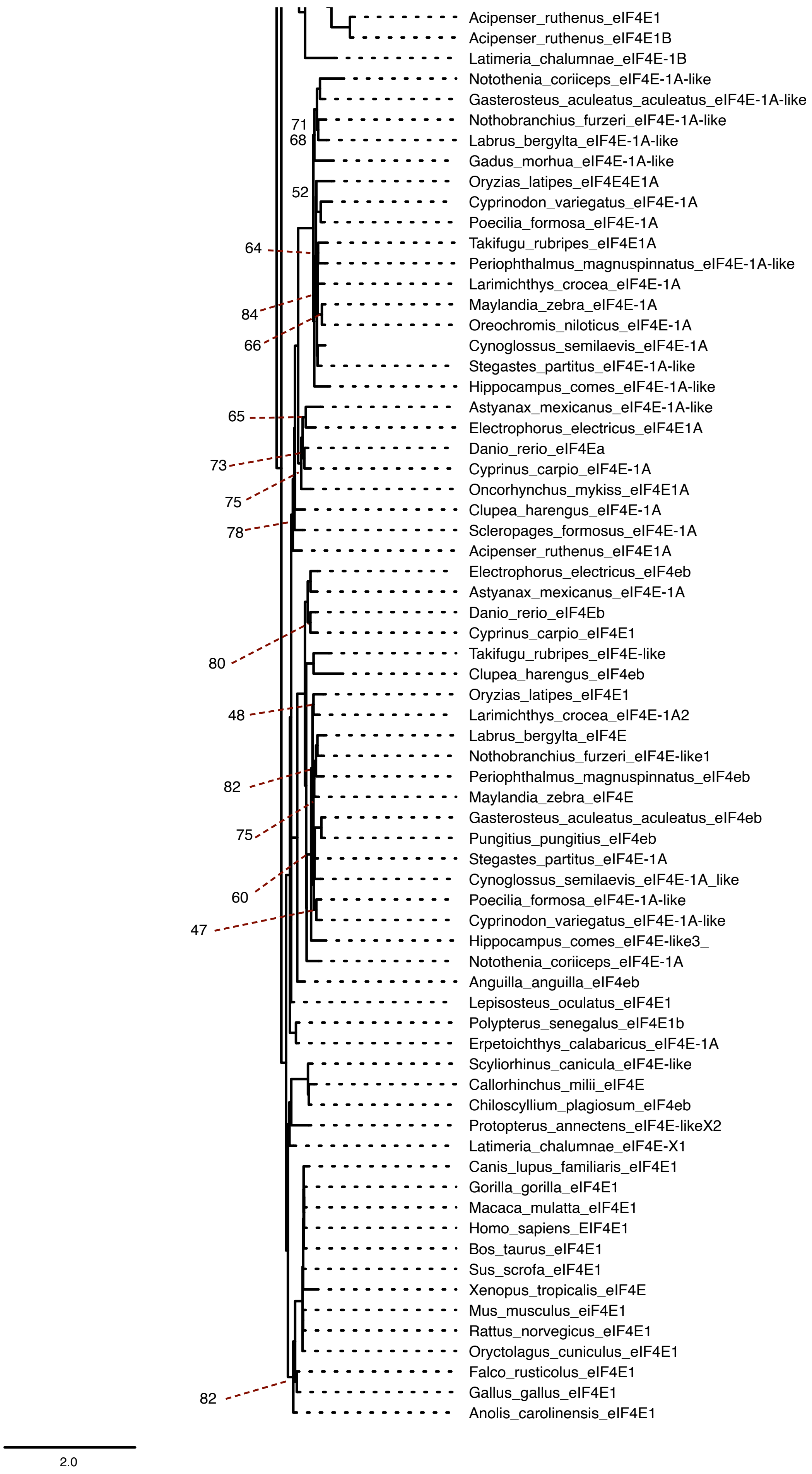

### Figure S1B

```
Callorhinchus      NRWALWYFKNDKTKSWTENLRLIAKFDTVEDFWALYNHIQQPSKLLFGCDYCLFKDGIKP 103
Scyliorhinus       NRWALWYFKNDKTKGWTENLRLIAKFDTVEDFWALYNHIQQPSKLLFGCDYCLFKDGIKP 105
Chiloscyllium      NRWALWYFKNDKTKSWTENLRLIAKFDTVEDFWALYNHIQQPSKLLFGCDYCLFKDGIKP 105
Protopterus        NRWALWYFKNDKSKSWTENLRLIAKFDTVEDFWALYNHIQRPSKLQFGCDYCLFKDGIKP 99
Oncorhynchus       NKWALWYFKNDKSKSWTENLRLIAKFDTVEDFWALYNHIQQPSKLGFGCDYCLFKDGVKP 98
Oryzias            NRWALWYFKNDKTKSWTENLRLISKFDTVEDFWALYNHIQQPSKLVLGCDYCLFKDGIKP 94
Gadus              NKWALWYFKNDKSKSWTENLRLISKFDTVEDFWALYNHIQQPSKLGFGCDYCLFKDGIKP 93
Notothenia         NRWALWYFKNDKSKSWTENLRLISKFDTVEDFWALYNHIQQPSKLGFGCDYCLFKDGIKP 106
Anguilla           NRWALWYFKNDKSKSWTENLRLIAKFDTVEDFWALYNHIQQPSKLGYGCDYCLFKDGIKP 95
Gasterosteus       NRWALWYFKNDKSKSWTENLRLISKFDTVEDFWALYNHIQQPSKLGFGCDYSLFKDGIKP 92
Pungitius          NRWALWYFKNDKSKSWTENLRLISKFDTVEDFWALYNHIQQPSKLGYGCDYSLFKDGIKP 94
Nothobranchius     NRWALWYFKNDKSKSWTENLRLISKFDTVEDFWALYNHIQQPSKLNFGCDYCLFKDGIKP 93
Takifugu           NRWALWYFKNDKSKSWTENLRLISKFDTVEDFWALYNHIQQPSKLGFGCDYCLFKDGIKP 93
Labrus             NRWALWYFKNDKSKSWTENLRLISKFDTVEDFWALYNHIQQPSKLGFGCDYCLFKDGIKP 94
Cynoglossus        NKWALWYFKNDKSKSWTENLRLISKFDTVEDFWALYNHIQQPSKLGFGCDYCLFKDGIKP 91
Periophthalmus     NRWALWYFKNDKSKSWTENLRLISKFDTVEDFWALYNHIQQPSKLGFGCDYCLFKDGIKP 93
Poecilia           NRWALWYFKNDKSKSWTENLRLISKFDTVEDFWALYNHIQQPSKLGFGCDYSLFKDGIKP 93
Cyprinodon         NRWALWYFKNDKSKSWTENLRLISKFDTVEDFWALYNHIQQPSKLGYGCDYSLFKDGIKP 93
Stegastes          NRWALWYFKNDKSKSWTENLRLISKFDTVEDFWALYNHIQQPSKLGFGCDYCLFKDGIKP 93
Maylandia          NRWALWYFKNDKSKSWTENLRLISKFDTVEDFWALYNHIQQPSKLGFGCDYCLFKDGIKP 93
Oreochromis        NRWALWYFKNDKSKSWTENLRLISKFDTVEDFWALYNHIQQPSKLGFGCDYCLFKDGIKP 93
Larimichthys       NRWALWYFKNDKSKSWTENLRLISKFDTVEDFWALYNHIQQPSKLGFGCDYCLFKDGIKP 93
Latimeria          NKWALWYFKNDKSKSWTENLRLIAKFDTVEDFWALYNHIQQPSKLQFGCDYCLFKDGIKP 116
Clupea             NRWALWYFKNDKSKSWTENLRLIAKFDTVEDFWALYNHIQQPSKLGFGCDYCLFKDGIKP 117
Danio              NRWALWYFKNDKSKSWTENLRLISKFDTVEDFWALYNHIQQPSKLGFGCDYCLFKDGIKP 96
Cyprinus           NRWALWYFKNDKSKSWTENLRLISKFDTVEDFWALYNHIQQPSKLGFGCDYSLFKDGIKP 96
Astyanax           NRWALWYFKNDKSKSWTENLRLISKFDTVEDFWALYNHIQQPSKLGFGCDYCLFKDGIKP 114
Electrophorus      NRWALWYFKNDKSKSWTENLRLISKFDTVEDFWALYNHIQQPSKLGFGCDYCLFKDGIKP 95
Scleropages        NRWALWYFKNDKSKSWSENLRLIAKFDTVEDFWALYNHIQQPSKLGFGCDYCLFKDGIKP 95
Acipenser          NKWALWYFKNDKSKSWTENLRLIAKFDTVEDFWALYNHIQQPSKLGFGCDYCLFKDGIKP 95
Lepisosteus        NRWALWYFKNDKSKSWTENLRLIAKFDTVEDFWALYNHIQQPSKLGFGCDYCLFKDGIKP 95
Polypterus         NRWALWYFKNDKSKSWTENLRLIAKFDTVEDFWALYNHIQQPSKLGFGCDYCLFKDGIKP 95
Erpetoichthys      NRWALWYFKNDKSKSWTENLRLIAKFDTVEDFWALYNHIQQPSKLGFGCDYCLFKDGIKP 96
Hippocampus        NRWALWYFKNDKSKSWTENLRLISKFDTVEDFWALYNHIQQPSKLVYGCDYCLFKDGIKP 100
Petromyzon         NRWALWFYKNDKSKSWQANLRLITKVDTVEDFWALYNHIQVASRLMPGCDYSLFKDGIEP 102
                   *:****::****:* *  *****:*.**************  *.*  ****.*****::*
```

Sharks

Lungfish

Seahorse

Lamprey

Spotted Gar – recently diverged from teleost

Gray bichir – dragon eel – similar to lungfish

Mudskippers – technically teleost but use fins as legs

Reedfish – bichir – snake fish

Eel

Teleost

### Figure S1B continued

```
Callorhinchus    MWEDDKNKKGGRWLMTLTKQQRHNDLDRYWLETLLCLIGEAFDEHSDEVCGAVVNVRPKG 163
Scyliorhinus     MWEDDKNKKGGRWLMTLNKQQRHNDLDRYWLETLLCLIGEAFDEHSDEVCGAVVNVRPKG 165
Chiloscyllium    MWEDDKNKRGGRWLMTLNKQQRHNDLDRYWLETLLCLIGEAFDEFSDEVCGAVVNVRPKG 165
Protopterus      MWEDELNKQGGRWLITLNKQQRHNDLDRYWLETLLCLIGEAFNDYSDDVCGAVVNVRPKG 159
Oncorhynchus     MWEDDKNKLGGRWLMTLSKQQRQIDLDRYWMETLLCLIGESFDEASEDVCGAVVNVRPKG 158
Oryzias          MWEDDKNKLGGRWLMTLNKQ-KHNDLDRYWMETLLCLVGESFDDASEEVCGAVVNVRHKG 153
Gadus            MWEDDRNKLGGRWLMTLNKQQRHNDLDRYWMETLLCLVGESFDESSEDVCGAVVNVRPKG 153
Notothenia       MWEDDRNKLGGRWLMTLNKQQRHNDLDRYWMETLLCLVGESFDAASEDVNGAVVNVRPKG 166
Anguilla         MWEDDRNKLGGRWLMTLNKQQRHNDLDRYWMETLLCLIGESFDEASDDVCGAVVNVRPKG 155
Gasterosteus     MWEDDRNKLGGRWLMTLNKQQRHNDLDRYWMETLLCLVGESFDEASEDVCGAVVNVRPKG 152
Pungitius        MWEDDRNKLGGRWLMTLNKQQRHNDLDRYWMETLLCLVGESFDEASEDVCGAVVNVRPKG 154
Nothobranchius   MWEDERNKLGGRWLITLNRQQRHNDLDRYWMETLLCLVGESFDEASDDVCGAVVNVRPKG 153
Takifugu         MWEDDRNKLGGRWLMTLNKQQRHNDLDRFWMETLLCLVGESFDEASDDVCGAVVNVRPKG 153
Labrus           MWEDDRNKLGGRWLMTLNKQQRHNDLDRYWMETLLCLVGESFDEASEDVCGAVVNVRPKG 154
Cynoglossus      MWEDDRNKLGGRWLMTLNKQQRHNDLDRYWMETLLCLVGESFDEASEDVCGAVVNVRPKG 151
Periophthalmus   MWEDDRNKLGGRWLMTLNKQQRHNDLDRYWMETLLCLVGESFDEASEDVCGAVVNVRPKG 153
Poecilia         MWEDDRNKLGGRWLMTLNKQQRHNDLDRYWMETLLCLVGESFDEASEEVCGAVVNVRPKG 153
Cyprinodon       MWEDDRNKLGGRWLMTLNKQQRHNDLDRYWMETLLCLVGDSFDEASEDVCGAVVNVRPKG 153
Stegastes        MWEDDRNKLGGRWLMTLNKQQRHNDLDRYWMETLLCLVGESFDEASEDVCGAVVNVRPKG 153
Maylandia        MWEDDRNKLGGRWLMTLNKQQRHNDLDRYWMETLLCLVGESFDEASEDVCGAVVNVRPKG 153
Oreochromis      MWEDDRNKLGGRWLMTLNKQQRHNDLDRYWMETLLCLVGESFDEASEDVCGAVVNVRPKG 153
Larimichthys     MWEDDRNKLGGRWLMTLNKQQRHNDLDRYWMETLLCLVGESFDEASEDVCGAVVNVRPKG 153
Latimeria        MWEDENNKRGGRWLMTLNKQQRHNDLDRYWLETLLCLIGESFDEHSDDVCGAVVNVRPKG 176
Clupea           MWEDDRNKLGGRWLMTLSKQQRHNDLDRYWMETLLCLIGESFDEASEDACGAVVNVRPKG 177
Danio            MWEDDRNKLGGRWLMTLSKQQRHNDLDRYWMETLLCLIGESFDEASEDVCGAVVNVRPKG 156
Cyprinus         MWEDDRNKLGGRWLMTLSKQQRHNDLDRYWMETLLCLIGESFDEASEDVCGAVVNVRPKG 156
Astyanax         MWEDDRNKLGGRWLMTLSKQQRHNDLDRYWMETLLCLIGESFDEASEDVCGAVVNVRPKG 174
Electrophorus    MWEDDRNKLGGRWLMTLNKQQRHNDLDRYWMETLLCLIGESFDEASEDVCGAVVNVRPKG 155
Scleropages      MWEDDRNKLGGRWLITLSKQQRHNDLDRYWMETLLCLIGESFDDASEDICGAVVNVRPKG 155
Acipenser        MWEDDRNKLGGRWLMTLNKQQRHNDLDRYWMETLLCLIGESFDEASEDVCGAVVNVRPKG 155
Lepisosteus      MWEDDRNKLGGRWLMTLGKQQRHNDLDRYWMETLLCLIGESFDEASDDVCGAVVNVRPKG 155
Polypterus       MWEDDRNKLGGRWLITLSKQQRHNDLDRYWMETLLCLIGESFDEASEDVCGAVVNVRPKG 155
Erpetoichthys    MWEDDRNKLGGRWLITLSKQQRHNDLDRYWMETLLCLIGESFDEASEDVCGAVVNVRPKG 156
Hippocampus      MWEDDRNKLGGRWLMTLNKVQRHNDLDRYWMETLLCLVGESFDEASDDVCGAVVNVRHKA 160
Petromyzon       MWEDERNKRGGRWLITLTKTQRHSDLDRYWLETLLCLIGEAFDDHSDDVCGAVVNVRPKA 162
                 ****: *: *****:** :  :: ****:*:******:*::*:  *::  ***:*** *.
```

**Sharks**

**Lungfish**

**Seahorse**

**Lamprey**

**Spotted Gar – diverged from teleost**

**Gray bichir – dragon eel – appears lungfish**

**Mudskippers – technically teleost but use fins as legs**

**Reedfish – bichir – snake fish**

**Eel**

**Teleost**

Figure S1B continued

```
Callorhinchus      DKISIWTGNCQSREAVTSIGQSYKERLGLPMKALIGYQSHDDTSSKSGSTTKNLYTV   220
Scyliorhinus       DKIAIWTGNCQNREAIVSIGQLYKERLGLSLKALIGYQSHDDTSSKSGSTTKNLFSV   222
Chiloscyllium      DKIAIWTGNCQNRDAVVSIGQLYKERLGLSLKALIGYQSHDDTSSKSGSTTKNLFSV   222
Protopterus        DKLSIWTTNCQNREAVVSIGQSYKERLGLPLKPVIGYQSHDDTSTKSGSTTKNLYSA   216
Oncorhynchus       DKISIWTGNCQNKEAIVAIGQQYKERLSIPIKLLIGYQSHDDTSSKSGSTTKNMYSV   215
Oryzias            DKISIWTGNCQNKEAIMTIGQLYKERLNLPMKAIIGYQSHDDTSSKSGSTTKNMYSV   210
Gadus              DKIAIWTSNCQNRDAIVTIGAGYKERLCLPSKPLISYQSHDDTSSKSGSTTKNMYSV   210
Notothenia         DKISIWTSNCQNRDAIMTIGQNYKERLSIPTKAIIGYQSHDDTSSKSGSTTKNIFSV   223
Anguilla           DKLAIWTGNCQNRDAIMTIGQQYKERLNIPSKALIGYQSHDDTSSKSGSTTKNMYTV   212
Gasterosteus       DKISIWTSQCQNRDAIMTIGQNYKERLNIPTKAIIGYQSHDDTSSKSGSTTKNMYSV   209
Pungitius          DKISIWTSQCQNRDAIMTIGQNYKERLNIPTKAIIGYQSHDDTSSKSGSTTKNMYSV   211
Nothobranchius     DKLAIWTSNCQNREAIMTIGQLYKERLSLPVKALIGYQSHDDTSSKSGSTTKNMYSV   210
Takifugu           DKIAIWTSNCQNREAIMTIGQLYKERLNIPIKAMLGYQSHDDTSSKSGSTTKNMYSI   210
Labrus             DKISIWTSNCQNRDAIMTIGQLYKERLTVPIKALIGYQSHDDTSSKSGSTTKNMYSV   211
Cynoglossus        DKIAIWTSNCQNREAIMTIGQQYKERLNIPIKAMIGYQSHDDTSSKSGSTTKNMYSV   208
Periophthalmus     DKIAIWTSNCQNRDAIMTIGQLYKERLSIPMKALIGYQSHDDTSSKSGSTTKNMYSV   210
Poecilia           DKISIWTSNCQNREAIMTIGQLYKERLNIPIKAMIGYQSHDDTSSKSGSTTKNMYSV   210
Cyprinodon         DKISIWTSNCQNREAIMTIGQLYKERLNLPIKAMIGYQSHDDTSSKSGSTTKNMYSV   210
Stegastes          DKIAIWTSNCQNREAIMTIGQLYKERLNLPIKAMIGYQSHDDTSSKSGSTTKNMYSV   210
Maylandia          DKISIWTSNCQNRDAIMTIGQLYKERLNLPMKAMIGYQSHDDTSSKSGSTTKNMYSV   210
Oreochromis        DKISIWTSNCQNRDAIMTIGQLYKERLNLPMKAMIGYQSHDDTSSKSGSTTKNMYSV   210
Larimichthys       DKIAIWTSNCQNRDAIMTIGQLYKERLNIPIKAMIGYQSHDDTSSKSGSTTKNMYSV   210
Latimeria          DKIAIWTTSCQNREAIMSIGQSYKERLGLPLKALIGYQSHDDTSSKSGSTTKNMYTV   233
Clupea             DKIAIWTANCQNRESIMTIGQQYKERLSIPNKTLIGYQSHDDTSSKSGSTTKNMYSV   234
Danio              DKIAIWTGNCQNRDAIMTIGQQYKERLSLPSKTLIGYQSHDDTSSKSGSTTKNMYSV   213
Cyprinus           DKISIWTGNCQNRDAIMTIGQQYKERLSLPIKTLIGYQSHDDTSSKSGSTTKNMYSV   213
Astyanax           DKIAIWTGNCQNRDAIMTIGLQYKERLNLPIKTLIGYQSHDDTSSKSGSTTKNMYSV   231
Electrophorus      DKIAIWTCNCQNRDAIMTIGQQYKERLNLPIKTLIGYQSHDDTSSKSGSTTKNMYSV   212
Scleropages        DKIAIWTVNCQNREAIMTIGQQYKERLNVPVKSLIGYQSHDDTSSKSGSTTKNMYTV   212
Acipenser          DKLAIWTGNCENKEAIMTIGQQYKERLSVPSKALIGYQSHDDTSSKSGSTTKNIFTV   212
Lepisosteus        DKISIWTGNCQNKEAIMTIGQQYKERLNVPNKALLGYQSHDDTSSKSGSTTKNMYTV   212
Polypterus         DKIAIWTGNCQNKEAIMTIGQLYKERLNMPLKALLGYQSHDDTSSKSGSTTKNMYTV   212
Erpetoichthys      DKIAIWTGNCQNKEAIMTIGQLYKERLNMPLKAILGYQSHDDTSSKSGSTTKNMYTV   213
Hippocampus        DKIAIWTSNCQNREAIMKIGETYKDLLSIPSKAMLGYQSHDDTSSKSGSTTKNMYSI   217
Petromyzon         DKIAVWTADCDNRESVVGIGRVYKDRLALPPRIIIGYQSHTDTATKSGSSTKNMFTV   219
                   **:::** . :.::::  **  **:** :  : ::.**** **::****:*** :
```

**Sharks**

**Lungfish**

**Seahorse**

**Lamprey**

**Spotted Gar – diverged from teleost**

**Gray bichir – dragon eel – appears lungfish**

**Mudskippers – technically teleost but use fins as legs**

**Reedfish – bichir – snake fish**

**Eel**

**Teleost**

**Supp Fig. 1. Phylogeny and conservation of eif4E orthologs.**

(A) Shown is the phylogeny of Eif4e1c orthologues (all 53 species with 142 homologs) estimated using maximum likelihood using IQ-TREE. Nodes with bootstrap support < 0.85 are marked with their respective values, all other nodes had support values of 0.85 or higher. (B) ClustalW sequence alignment of Eif4e1c from aquatic vertebrate. Species names are color-coded by type with the legend on the bottom. Conserved Eif4e1c specific residues are highlighted in red bold with similar residues highlighted in pink.

### Figure S2

A

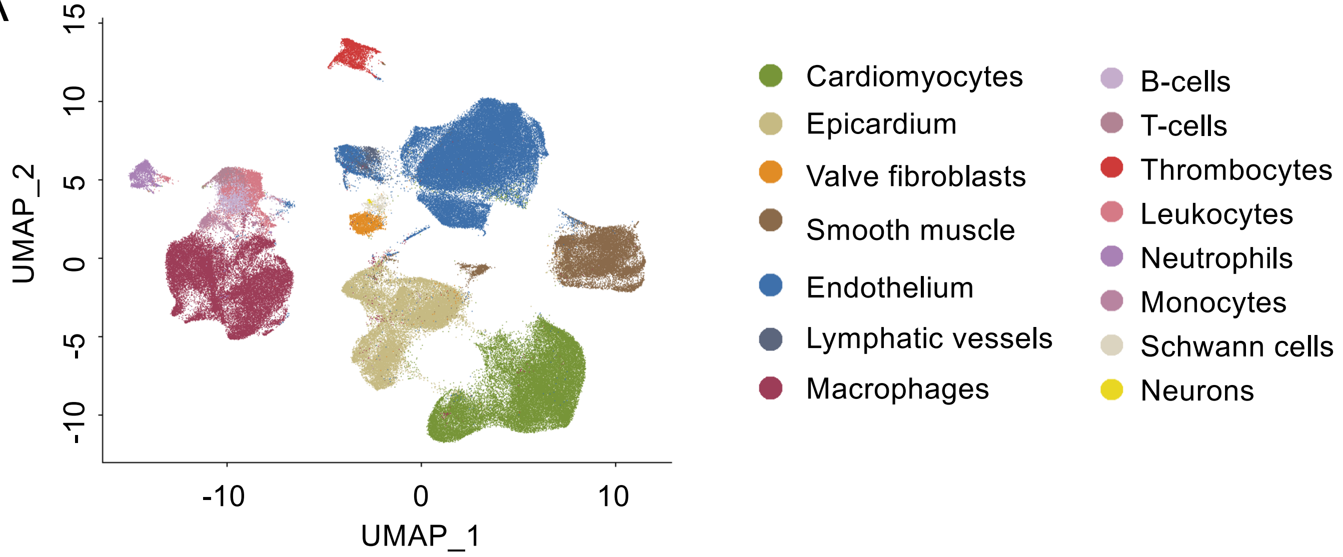


B

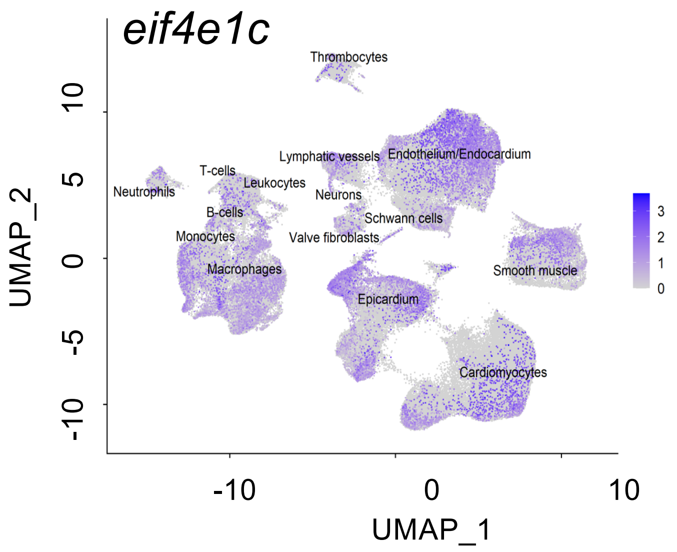


| Cell-types | *% eif4e1c* (+) cells | *% eif4ea* (+) cells | *% eif4eb* (+) cells |
|---|---|---|---|
| Cardiomyocytes | 4.58 | 1.45 | 4.47 |
| Epicardium | 17.51 | 7.98 | 14.46 |
| Valve fibroblasts | 7.49 | 3.39 | 10.97 |
| Smooth muscle | 9.56 | 5.50 | 9.86 |
| Endothelium | 10.64 | 4.44 | 7.27 |
| Lymphatic vessels | 12.86 | 4.36 | 8.18 |
| Macrophages | 13.67 | 4.53 | 5.36 |
| B-cells | 3.78 | 1.19 | 0.89 |
| T-cells | 1.94 | 0.91 | 0.67 |
| Thrombocytes | 2.69 | 0.96 | 1.69 |
| Leukocytes | 8.30 | 2.26 | 2.56 |
| Neutrophils | 5.89 | 2.26 | 2.39 |
| Monocytes | 11.49 | 6.42 | 3.54 |
| Schwann cells | 8.35 | 4.85 | 6.80 |
| Neurons | 42.86 | 14.29 | 2.04 |

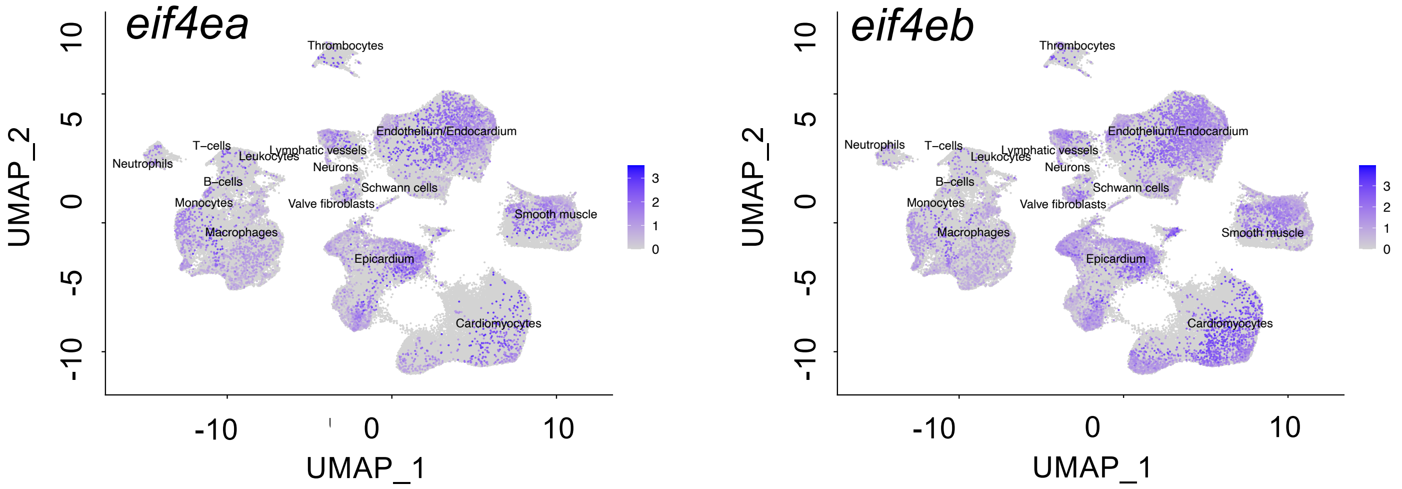

**Supp Fig. 2. EIF4E homologs are expressed in all cell-types within zebrafish hearts.** (A) UMAP representation of single-cell RNA-seq data and clustering results of zebrafish hearts. (B) UMAP plots depicting *eif4e1c*, *eif4ea* and *eif4eb* expression. Right - Table showing the distribution of *eif4e1c* expressing cells in each cell clusters.

Figure S3

A

| Cross | Stage | N | WT | HETS | HOMS | chi² | p value | Significant? |
|---|---|---|---|---|---|---|---|---|
| Δeif4e1c HETS X HETS | 3dpf | 190 | 51 | 85 | 54 | 1.94 | 0.33 | no |
| | 4wpf | 410 | 109 | 203 | 98 | 0.629 | 0.73 | no |
| | 8wpf | 630 | 207 | 312 | 111 | 29.3 | 4.31E-07 | yes |
| | >3mpf | 1868 | 599 | 951 | 318 | 84.9 | 3.22E-19 | yes |
| Δeif4e1c HOMS ♀ X HETS ♂ | 3dpf | 107 | N/A | 58 | 49 | 0.757 | 0.384 | no |

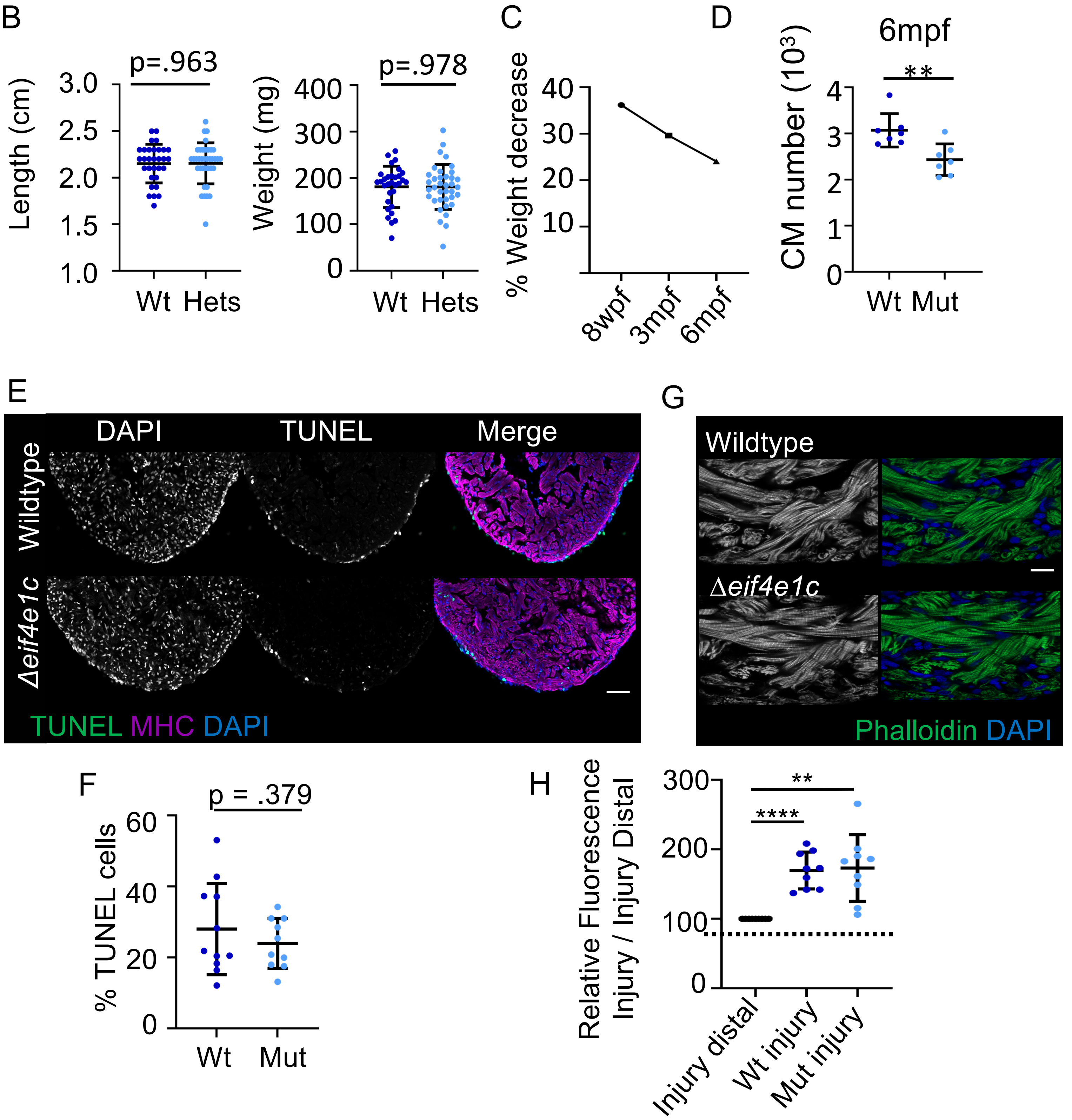

**Supp Fig. 3. Heterozygotes for D*eif4e1c* were compared for length and weight with their wildtype siblings.** (A) Table showing survival of progeny from crosses of Deif4e1c heterozygote carriers. Clutches of fish were genotyped at the listed development stage (dpf, wpf and mpf represent days-, weeks- or months-post-fertilization). Significant deviations from Mendelian ratios were calculated using $chi^2$ and the resulting p-value is shown. (B) Fish were grown together, genotyped at the listed stage (top label) and immediately measured from jaw to the caudal fin bifurcation (Average length WT= 2.15cm and mutant= 2.15cm with N = 31 vs 37; Welch's t-test, p-value =0.963,). After measuring length, fish were dried and weighed (Average weight: wildtype = 181.0mg and mutant = 180.7mg with N = 31 vs 37; Welch's t-test, p-value = 0.978,). (C) The percent weight decrease between mutant and wildtype fish was calculated for each of the time points (D) CM numbers for wildtype and D*eif4e1c* mutants were counted at 6mpf from cryosections stained with an antibody for Mef2c (Scale bar = 100mm). The numbers of Mef2c positive cells were counted with MIPAR (average wildtype = 3070 and mutant = 2432 with N = 7 vs 7; Welch's t-test, p-value = 0.0055). (D) TUNEL staining of zebrafish hearts (Scale bar = 50mm). (F) Quantification of TUNEL stain showed no increase in the number of cells undergoing apoptosis in adult hearts (mean: wildtype = 28.03, mutant = 23.98; Welch's t-test, p-value = .379, N = 11 vs 10) (G) Phalloidin staining of wildtype (top) and mutant (bottom) zebrafish hearts. Left – Phalloidin in gray scale. Right – Merge of phalloidin with DAPI (Scale bar = 10mm). (H) Quantification of Eif4ea/b immunostaining from Figure 3G (wildtype mean increase (blue) = 1.70, Mann-Whitney p-value < 0.0001, N = 9 vs 9; mutant mean increase (light blue) = 1.73, Mann-Whitney p-value = 0.0018, N = 9 vs 9). Horizontal black bars display the mean (middle) or standard error (top and bottom).

Figure S4

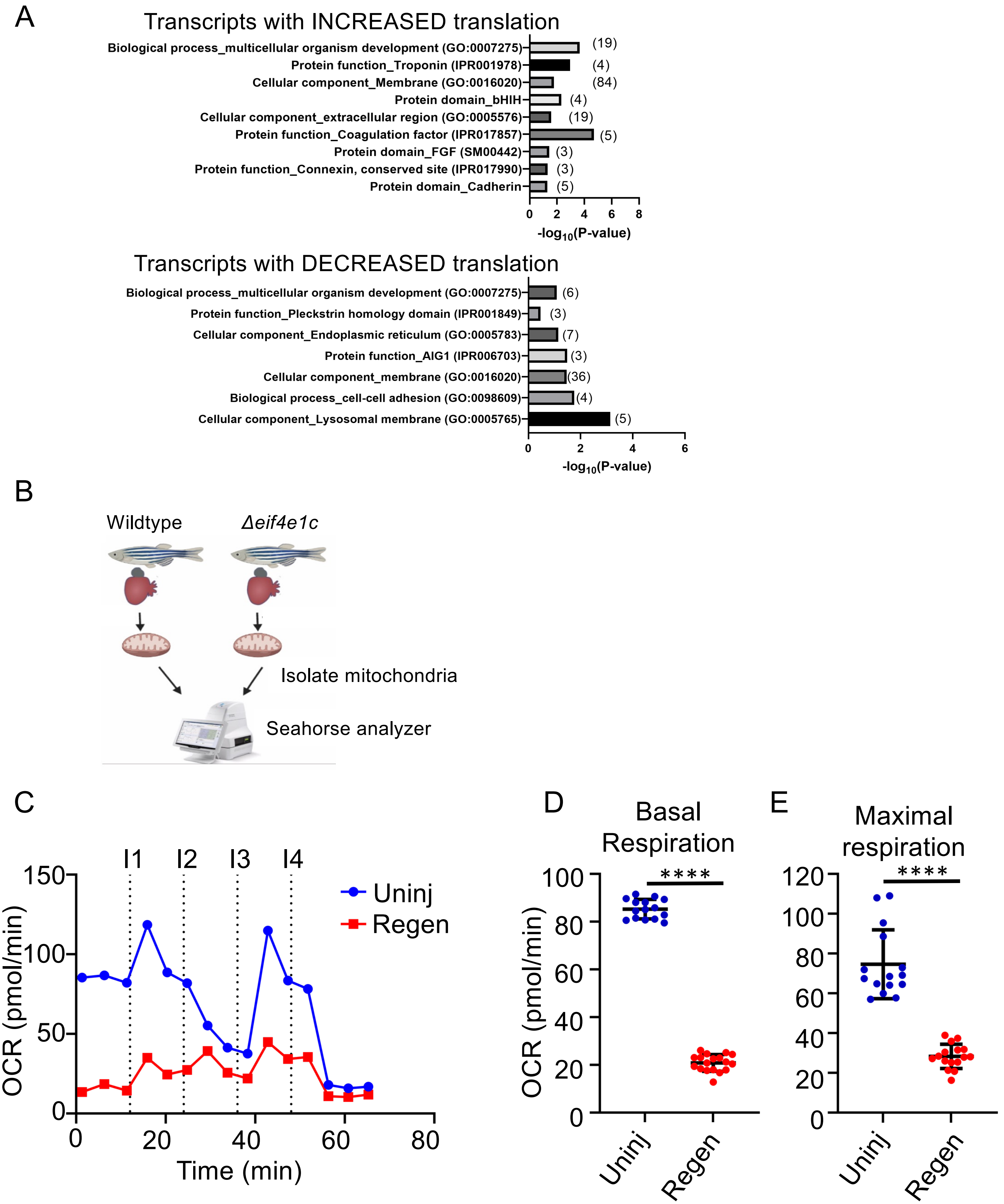
A
Transcripts with INCREASED translation
Biological process_multicellular organism development (GO:0007275) (19)
Protein function_Troponin (IPR001978) (4)
Cellular component_Membrane (GO:0016020) (84)
Protein domain_bHLH (4)
Cellular component_extracellular region (GO:0005576) (19)
Protein function_Coagulation factor (IPR017857) (5)
Protein domain_FGF (SM00442) (3)
Protein function_Connexin, conserved site (IPR017990) (3)
Protein domain_Cadherin (5)
-log10(P-value)
Transcripts with DECREASED translation
Biological process_multicellular organism development (GO:0007275) (6)
Protein function_Pleckstrin homology domain (IPR001849) (3)
Cellular component_Endoplasmic reticulum (GO:0005783) (7)
Protein function_AIG1 (IPR006703) (3)
Cellular component_membrane (GO:0016020) (36)
Biological process_cell-cell adhesion (GO:0098609) (4)
Cellular component_Lysosomal membrane (GO:0005765) (5)
-log10(P-value)
B
Wildtype
Δeif4e1c
Isolate mitochondria
Seahorse analyzer
C
I1 I2 I3 I4
Uninj
Regen
OCR (pmol/min)
Time (min)
D
Basal Respiration
OCR (pmol/min)
****
Uninj
Regen
E
Maximal respiration
****
Uninj
Regen

**Supp. Fig. 4.**

(A) Gene ontology categories were discovered using the DAVID on-line platform. Each category is listed on the left with the identifier in parentheses. Shown are GO categories with p-value < 0.05, x-axis is $-\log_{10}$(p-value). Parentheses to the right of the bar indicate numbers of genes identified in each group. (B) Cartoon depicting experimental set-up for Seahorse analysis. Mitochondria were isolated from hearts dissected from wildtype and mutant fish and then processed on the Seahorse analyzer from Agilent. (C) Shown is a time-course of oxygen consumption rates (OCR) from mitochondria measured by the Seahorse analyzer. Hashed lines indicate time points of drug injection. The first injection (I1) is of ADP to stimulate respiration, the second injection (I2) is of oligomycin to inhibit ATP synthase (complex V) decreasing electron flow through transport chain, the third injection (I3) is of FCCP to uncouple the proton gradient, and the fourth injection (I4) is of antimycin A to inhibit complex III to shut down mitochondrial respiration. (D) Basal respiration is calculated as the OCR average after addition of ADP (I1 to I2) subtracting the non-mitochondrial respiration after injection of antimycin A (after I4) (mean: wildtype uninjured= 85.30 and wildtype regenerating= 20.90; Welch's t-test p value < 0.0001; N= 15,18). (E) Maximal respiration is calculated as the OCR average after FCCP addition (I3 to I4) subtracting the non-mitochondrial respiration after injection of antimycin A (after I4) (mean: wildtype uninjured= 74.64 and wildtype regenerating= 28.34; Mann-Whitney p-value < 0.0001; N= 15,18). For panels C-E, individual data points are represented by dots, uninjured is in blue, regenerating is in red, and horizontal black bars display the mean (middle) or standard error (top and bottom). Wildtype – Blue, Mutant – light blue.

Figure S5

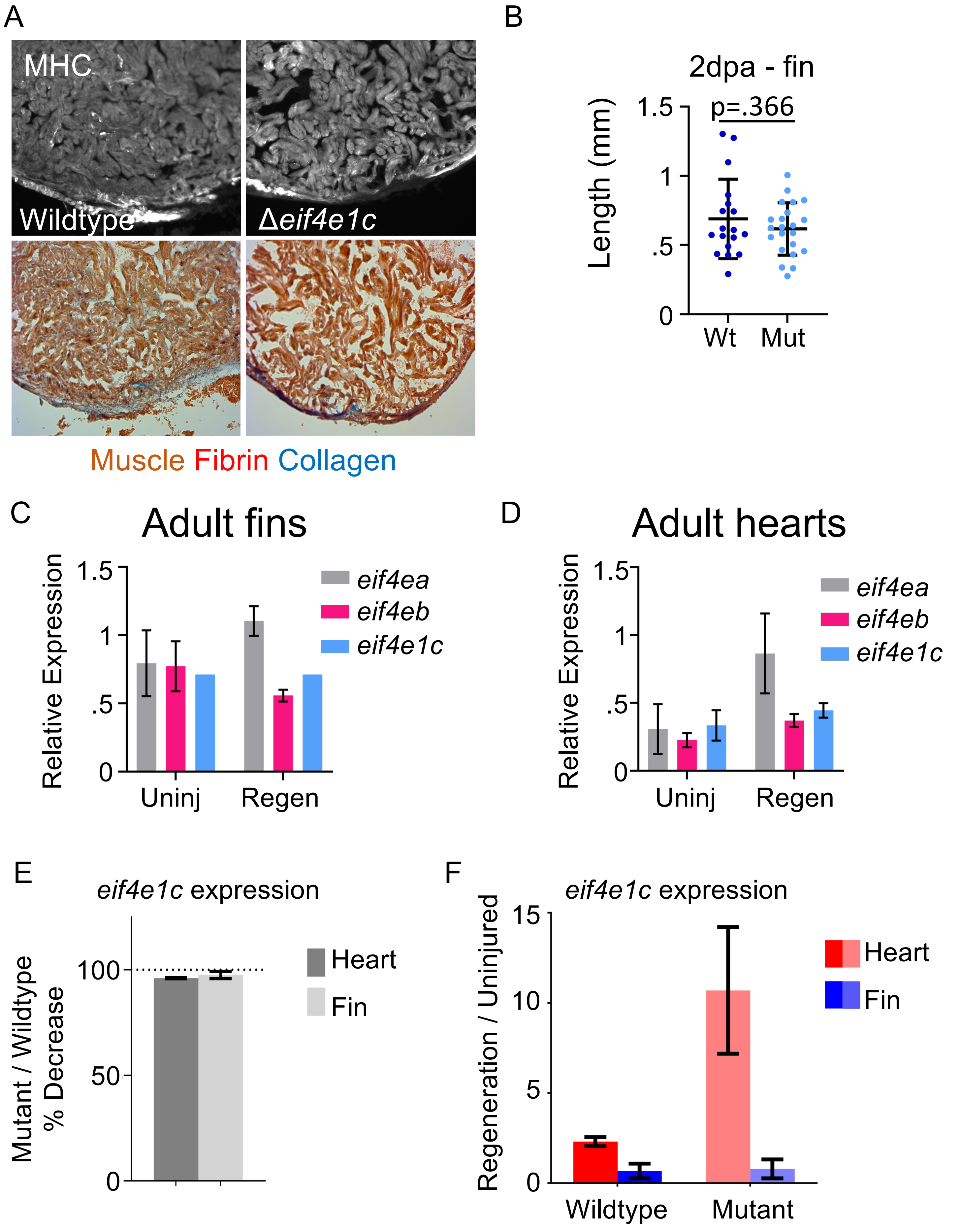
A
MHC
Wildtype
Δeif4e1c
Muscle Fibrin Collagen
B
2dpa - fin
Length (mm)
1.5
1
.5
0
p=.366
Wt
Mut
C
Adult fins
Relative Expression
1.5
1
.5
0
eif4ea
eif4eb
eif4e1c
Uninj
Regen
D
Adult hearts
Relative Expression
1.5
1
.5
0
eif4ea
eif4eb
eif4e1c
Uninj
Regen
E
eif4e1c expression
Mutant / Wildtype
% Decrease
100
50
0
Heart
Fin
F
eif4e1c expression
Regeneration / Uninjured
15
10
5
0
Heart
Fin
Wildtype
Mutant

**Supp. Fig. 5.**
(A) Images of sectioned ventricles at 28dpa are stained for Myosin heavy chain (MHC) to indicate cardiac muscle. Shown below are the same sections stained with AFOG which shows muscle (orange), Fibrin (red) and Collagen (blue). Hearts from 5 wildtype and 5 ∆*eif4e1c* mutants were ranked by injury size. Shown are representative hearts near the averages of the groups. (B) Caudal fins were amputated ~50% and the blastema were imaged 48 hours later for ∆*eif4e1c* mutants and their wildtype siblings. The length of the blastema was measured from the tip of the 3$^{rd}$ ray using the Zeiss microscope software. There was no significant difference between the groups (2dpa average wildtype = 0.689mm and mutant = 0.617mm; Welch's t-test, p-value = 0.366, N = 18 vs 22). Horizontal black bars display the mean (middle) or standard error (top and bottom). (C) Expression of each transcript was normalized to *mob4* in uninjured fins (left) and fins undergoing regeneration (right). (D) same as (C) but for the heart. (E) RT-qPCR analysis of *eif4e1c* from uninjured hearts (r dark gray) and fins (light gray) in ∆*eif4e1c* mutants. Ct values are normalized first to *mob4* and then to the wildtype values (hashed line). (F). RT-qPCR analysis of *eif4e1c* during regeneration of the heart (red) and the fin (blue). Wildtype is shown in dark colors (red and blue) and ∆*eif4e1c* mutants are shown in light colors (pink and light blue).

### Figure S6

A

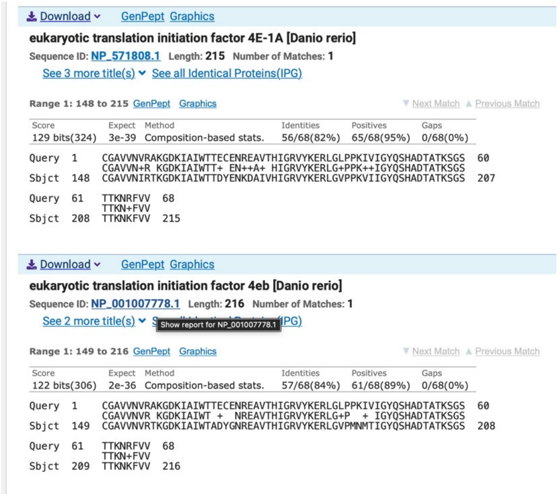


B

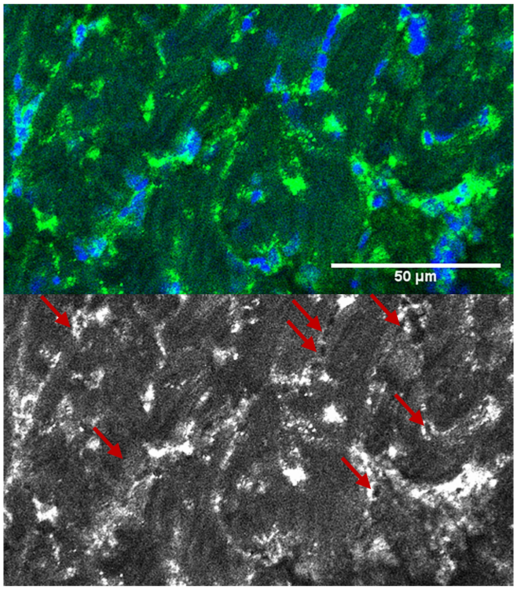


C

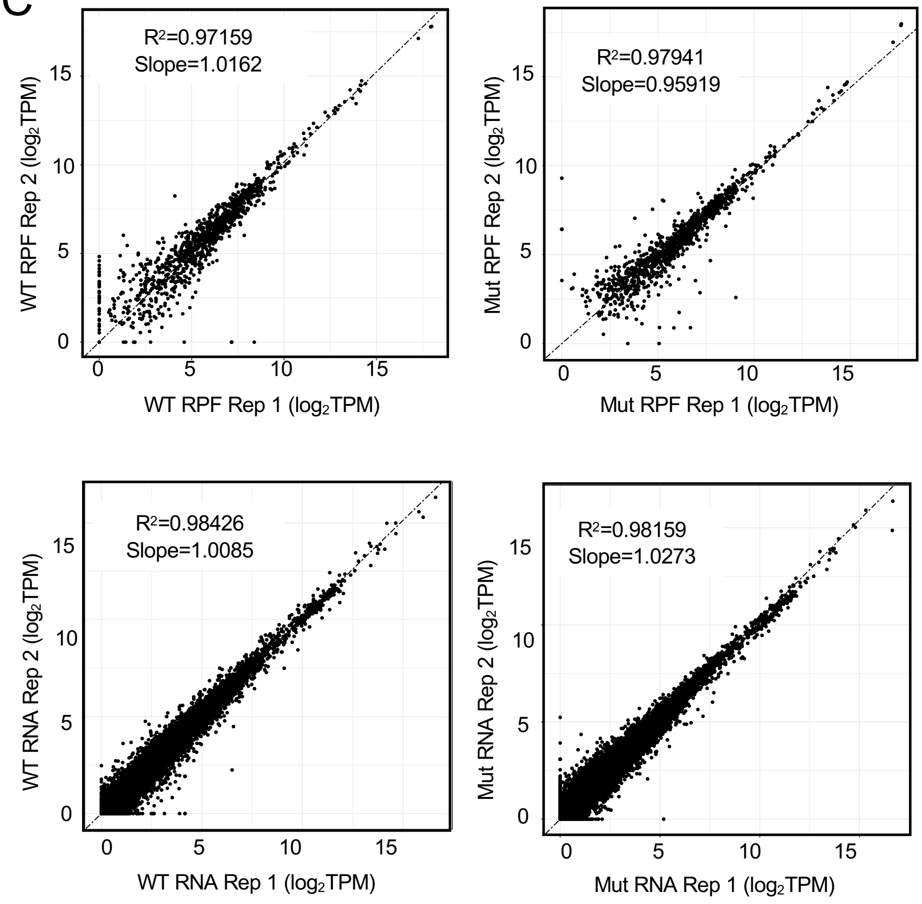

**Supp. Fig. 6.**

(A) Region of human EIF4E1 (query) used to raise ab33768 is nearly identical in both paralogs of zebrafish canonical Eif4e1 (subject). (B) Immunofluorescence staining with ab33768 (green) on zebrafish hearts produces the expected cytoplasmic staining pattern (nuclei – blue). (C) Shown are the correlations between replicates for ribosome profiling (top) and RNA sequencing of the inputs (bottom) for both wildtype (left) and ∆*eif4e1c* mutant (right) samples.
